# Go thou to the ant: A comparative biomechanical analysis of locomotion in Hymenoptera (Hexapoda)

**DOI:** 10.1101/2023.02.24.529971

**Authors:** V. Regeler, B.E. Boudinot, T. Wöhrl

**Affiliations:** Institut für Zoologie und Evolutionsforschung, Friedrich-Schiller-Universität Jena, 07743 Jena, Deutschland

**Keywords:** Functional morphology, evolution, cursoriality, kinematics, Formicidae.

## Abstract

Although ants are conceived of as paragons of social complexity, it may be their locomotory capacity that truly sets them apart from other Hymenoptera. Based on our comparative kinematic analysis of Formicidae for level, straight-line locomotion in a broad phylogenetic context, we observe that ants are distinctly capable runners. No sampled hymenopteran paralleled the body-scaled speed of ants. Relative stride lengths for ants were longer than other sampled taxa despite short ground contact durations relative to swing durations. With respect to spatial gait patterns, ants had relatively narrow hindleg and broad midleg step-widths on average, possibly enhancing speed and turning ability. Ants were able to extend their propulsive pair of legs, those of the metathorax, extremely far posterad relative to other sampled taxa, and had a distinct locomotory posture, with a high ground clearance and the femorotibial joints raised above their backs. Despite the unique modifications of their coxotrochanteral articulations, ant forelimbs were largely unremarkable with respect to our quantified variables. Sawflies, in contrast, had extremely wide and perhaps inefficient foreleg stances, and were observed for the first time to have what appears to be a dominant tetrapodal gait pattern, which raises unexpected questions about the early evolution of the Hymenoptera. Finally, we observed variability in attachment abilities and no consistent pattern of leg liftoff sequence across the sampled taxa. Our results establish locomotory evolution in the Hymenoptera as a functionally and structurally variable system with numerous directions of future research, particularly for phylogenetic comparison across wing-monomorphic and wing-polymorphic lineages.

**Summary statement:** This work establishes a comparative phylogenetic approach to hymenopteran kinematics, demonstrating that ant locomotory capacity is derived and observes, unexpectedly, that the sampled sawflies (symphyta) never used a tripod gait.

## 1. Introduction

The ants (Formicidae) are renowned for their industriousness, social complexity, and sheer diversity and abundance, comprising potentially 20,000 trillion individuals distributed across well over 14,000 described species (Schultheiss *et al.,* 2022; Bolton, 2023). Both covering and filling terrestrial surfaces globally, ants are model hexapodal locomotors for which diverse subjects have been investigated, including body scaling (Zollikofer, 1994b; Merienne *et al*., 2020; Tross *et al*., 2021, 2022), motion under load (Zollikofer, 1994c), incline traversal (Seidl & Wehner, 2008; Wöhrl *et al*., 2017, 2021), running mechanics (Zollikofer, 1994a; Reinhardt & Blickhan, 2014; Wahl *et al*., 2015; Pfeffer *et al*., 2019; Arroyave-Tobon *et al*., 2022), surface attachment (Federle *et al*., 2001; Endlein *et al*., 2008, 2013; Humeau *et al*., 2019), above and belowground navigation (Wittlinger *et al*., 2007; Gravish *et al*., 2013), saltation (Baroni Urbani *et al*., 1994; Ye *et al*., 2020), and swimming (Bohn *et al*., 2012; Yanoviak & Frederick, 2014; Schultheiss & Guénard, 2021). Remarkably, among the evolutionarily derived features of ants (*e.g.*, Wilson *et al.,* 1967a,b; Bolton, 2003; Boudinot, 2015; Boudinot *et al.,* 2022), it is only their proximal leg articulations that are developmentally and phylogenetically constrained between the winged males and wing-polyphenic females, the latter of which include the worker and queen castes. Specifically, the foreleg coxotrochanteral mechanism of Formicidae is uniquely monocondylic and the mid- and hindleg coxal bases are constricted into small hemispherical balls that fit into their corresponding thoracic sockets (Boudinot, 2015). Despite these derivations and the intense interest in their biomechanics, the functional consequences of the leg modifications of ants have never been investigated, nor has the locomotion of ants been explicitly studied in a comparative context among their relatives in the Hymenoptera.

The Hymenoptera themselves are one of the four “megadiverse” insect orders alongside Lepidoptera, Diptera, and Coleoptera (Grimaldi & Engel, 2005), and are ancestrally monomorphic for the occurrence of wings (Königsmann, 1976; Gauld & Bolton, 1988; Beutel *et al*., 2014; Hanna & Abouheif, 2021). Morphologically, they display extreme disparity of form (*e.g.*, Goulet & Huber, 1993) and span orders of magnitude for body length, with some individuals ranging from little over 100 microns (*Dicopomorpha echmepterygis*, 140 mm; Mockford, 1997) to nearly a decameter (*Pelecinus polyturator*, 90 mm; Johnson & Musetti, 1999). The diversification of crown-clade or modern-day Hymenoptera may have begun in the Permian or Triassic, with the broad-waisted, phytophagous, and paraphyletic “symphyta” (sawflies) occurring in the fossil record prior to the narrow-waisted and carnivorous Apocrita, followed eventually by the stinging wasps, Aculeata, from which the ants evolved (Rasnitsyn, 1975; Ronquist *et al.,* 1999, 2012; Vilhelmsen, 2009b; Peters *et al.,* 2017; Branstetter *et al.,* 2017; Boudinot *et al.,* 2022). Although winglessness in females has evolved numerous times throughout the Hymenoptera (*e.g.*, Hanna & Abouheif, 2021)—representing a major locomotory transition from the sky to the ground—studies of walking or running mechanics are restricted to a few species of ants (*e.g.*, Zollikofer, 1994a; Reinhardt & Blickhan, 2014; Wahl *et al.,* 2015) and the honey bee (Zhao *et al.,* 2018).

Numerous kinematic parameters have been established for the quantification and analysis of terrestrial locomotion in insects, some of which differ from those of bi- and quadrupedal vertebrates due to the distinct spatial and temporal dynamics of six-legged motion (*e.g.*, Hughes, 1952; Alexander, 2003; Wosnitza *et al.,* 2013; Wahl *et al.,* 2015; Wöhrl *et al.,* 2017; Pfeffer *et al.,* 2019). The so-called tripod gait of insects is characterized by the simultaneous work of the front and rear legs on one side of the body and of the middle leg on the other side (Hughes, 1952; Zollikofer, 1994a; Wendler, 1964). When the two groups of legs involved in the left and right tripod alternate one after the other, the pattern is referred to as the alternating tripod gait, which is typical of insects, and has been kinematically documented to some degree in apterygote insects (Manton, 1972; Tichy, 1988), cockroaches (Hughes, 1952; Full & Tu, 1991; Bender *et al.,* 2011), stick insects (Wendler, 1964; Graham, 1972;), grasshoppers and crickets (Whitney & Hedwig, 2011; Zhang *et al.,* 2015; Barreto *et al.,* 2021; Zhang *et al.,* 2022), leaf-footed bugs (Frantsevich & Cruse, 2005), beetles (Hughes, 1952, Zurek *et al.,* 2015), Lepidoptera (Johnston & Levine, 1996), and fruit flies (Strauß & Heisenberg, 1990; Wosnitza *et al.,* 2013), in addition to ants and honey bees. The tripod gait has been computationally shown to be favored for fast locomotion on three-dimensional terrain with sticky feet (Ramdya *et al*., 2017). Insects can move in other, sometimes more complex stepping patterns such as a tetrapod gait or a “wave gait” (Hughes, 1952; Grabowska *et al.,* 2012; Wosnitza *et al.,* 2013; Ambe *et al.,* 2018), or in unusual bipedal gaits when tarsal adhesion is perturbed (Ramdya *et al*., 2017).

Temporally, gait patterns are determined by the duration of the swing and contact phases across multiple steps. A swing phase is understood as the length of time between tarsal lift-off and touchdown, while the contact (or stance) phase is defined by the time between tarsal touchdown and lift-off (Wahl *et al.,* 2015; Reinhardt & Blickhan, 2014). Together, one contact and one swing phase comprise a full step cycle. The step frequency indicates the total number of step cycles that occur in one second (Hz; Reinhardt & Blickhan, 2014). The temporal step pattern can also be used to measure the swing-phase synchronicity of the legs involved in the tripod via the tripod coordination strength variable (TCS; Wosnitza *et al.,* 2013; Wahl *et al.,* 2015; Pfeffer *et al.,* 2019; Merienne *et al.,* 2020). Another measure of the step cycle is the duty factor, which quantifies the percentage of a total cycle that a given leg is in the contact phase (Weihmann *et al.,* 2017; Wahl *et al.,* 2015; Humeau *et al.,* 2019). The duty factor can, therefore, be taken as a measure to describe the transition from walking to running or of running itself (Alexander, 2003; Wahl *et al.,* 2015; Pfeffer *et al.,* 2019). According to Alexander (2003), one may refer to a running rather than a walking pace once the duty factor is below 50%, in which case the ground contact time is shorter than the swing phase. Insects usually have duty factors that are above the critical value of 50%, such that when the tripod is changed, all legs are in a ground contact phase for a brief moment and there are no pure swing phases, resulting in grounded or Groucho running, or compliant walking (Alexander & Jayes 1978; McMahon *et al*., 1987; Ting *et al.,* 1994; Rubenson *et al*., 2004; Reinhardt & Blickhan, 2014). There are exceptions to this generalization, however, such as the desert ant, *Cataglyphis fortis*, which has been shown to jump from step to step at a high speed of 369 mm/s, thus increasing its stride while consequently decreasing the duty factor all of their legs to below 50% (Wahl *et al.,* 2015; Pfeffer *et al.,* 2019).

Other kinematic parameters are also largely proportional to running speed, such as stride length (Wahl *et al.,* 2015; Wosnitza *et al.,* 2013; Nirody, 2021), which describes the distance covered by a tarsus from the time of one touchdown to the next (Wahl *et al.,* 2015; Pfeffer *et al.,* 2019; Alexander, 2003) and may be visualized via spatial stride pattern (Wahl *et al.,* 2015; Pfeffer *et al.,* 2019; Tross *et al.,* 2021). Spatial step patterns contain further information about the track or step width of the gait pattern, *i.e.*, the maximum left-to-right distance between the tarsal tips, which may inform questions of static and/or dynamic stability (*e.g.*, Ting *et al.,* 1994; Reinhardt, 2014). Finally, it is worthwhile to determine the range of motion for each leg for the spatial analysis of gait patterns. In this context, the anterior and posterior extreme positions (AEP, PEP) are defined as the tarsal position of touchdown and lift-off, respectively (Tross *et al.,* 2022; Cruse *et al.,* 2006; Seidl & Wehner, 2008). An AEP value of < 90° indicates tarsal touchdown anterad the coxa and > 90° posterad; the converse applies to the maximum angle or the liftoff position (PEP) of the tarsus. These positions can be used to determine the total range of motion of the legs during the run (Cruse, 1976; Merienne *et al.,* 2020) and can be used to visualize leg track trajectories (Bohn *et al.,* 2012; Tross *et al.,* 2022).

The present work is motivated by the central question: *Are ants different than other Hymenoptera in their locomotory capacities*? Although Formicidae are known to be defined by their leg joints and numerous studies have quantified their locomotion, no kinematic study exists to date that compares the patterns of ant motion to their relatives, be they close or distant. Our expectations at the outset of this work were that: (1) the predominant gait pattern would be tripodal; (2) ants and other Hymenoptera would be able to locomote at comparable speeds; and (3) the fore-, mid-, and hindlegs of ants will have a quantifiably distinct range of motion. To address our overarching question and to evaluate our specific expectations, we collected a phylogenetically diverse sample of Hymenoptera and quantified temporal and spatial locomotory data from a series of flat-surface, straight-line kinematic recordings. Using body length to control for size, our results clearly show differences between ants and other wasps, setting the foundation for future comparative-phylogenetic study of locomotion in Hymenoptera and other insects.

## 2. Materials and Methods

### 2.1. Materials

The tabular overviews of the materials, devices and software programs used, including their manufacturer information, as well as greater methodological detail can be found in the supplementary data. The supplementary tables include the materials involved in the camera construction and the composition of the lens (Tab. S1), the various components of the measuring range (Tab. S2), the materials and devices used for collecting and weighing (Tab. S3), and a list of the software programs used (Tab. S4).

### 2.2. Hymenoptera sampling

Specimens used in the present study were collected during August 2022 in Jena, Germany. The Hymenoptera were netted from the air, resting places, or food sources. After netting, specimens were carefully transferred to moisture chambers (25 mL centrifuge tubes), which were kept in a darkened place until the insect could be subject to kinematic filming. The specimens were identified by sight in the field, with later determinations made using a stereomicroscope and the guides of Goulet & Huber (1993) and Klausnitzer (2011). To render the most robust comparisons between ants and non-ants, and to link mechanics and evolution, our sampling strategy was explicitly phylogenetic: We aimed to span as many nodes in the hymenopteran phylogeny as possible from the local fauna, *i.e.*, to collect a maximally diverse sample given the known relationships within the order (Peters *et al.,* 2017; Branstetter *et al.,* 2017). In total, the movement of 38 individuals was recorded in 155 separate measurements, of which 28 individuals representing 21 genera and 14 families were selected for further analysis (Tables 2, S5). We sampled 10 surface foraging Formicidae belonging to four genera (*Camponotus*, *Formica*, *Lasius*, *Myrmica*) across two subfamilies (Formicinae, Myrmicinae). One sampled *Lasius* was a queen (i27), and one was a parasite infected worker (i35); we kept both specimens in the dataset as they did not affect our main conclusions and as their comparative performances were of interest. Our superfamilial nomenclature follows Pilgrim *et al*. (2008).

**Box 1.**
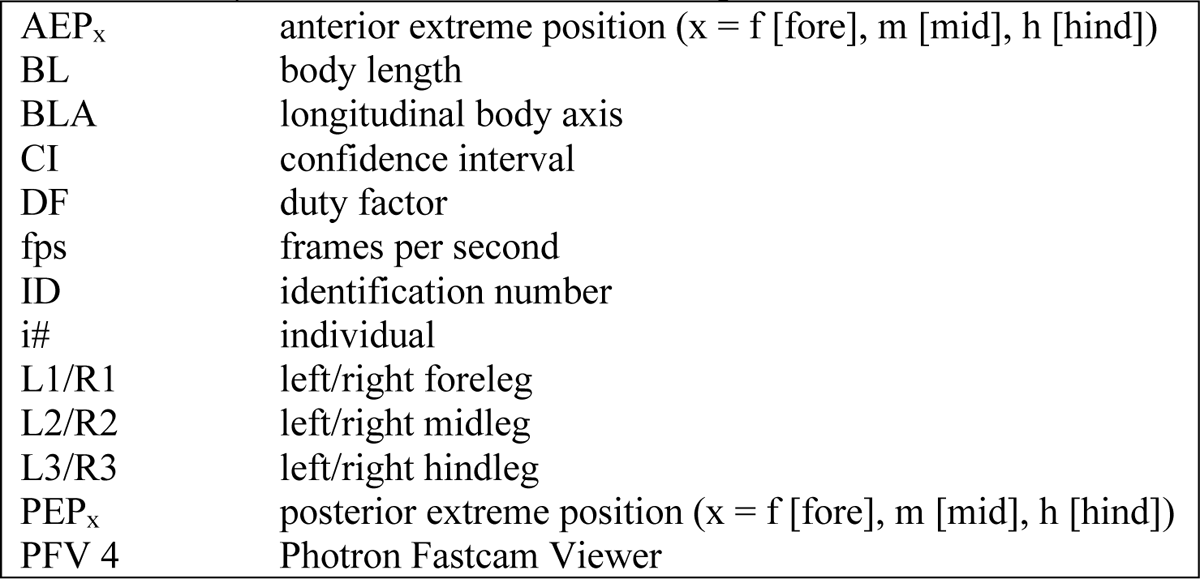
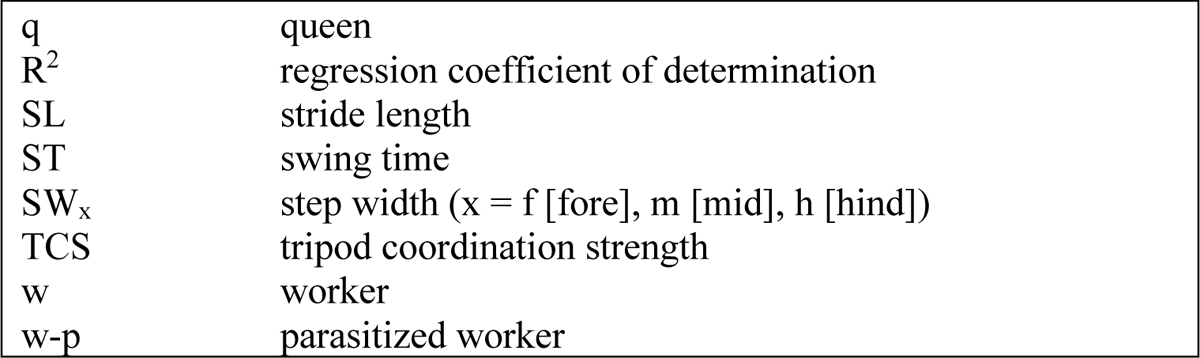
Summary of abbreviations used in the present work.

### 2.3. Experimental setup

For kinematic measurements, a high-speed Photron Fastcam SA3 (San Diego, CA, USA) camera with frequency of 500 Hz and equipped with a Componon-S enlarging lens, a C-mount lens adapter, and a Makro-Unifoc 12 focusing tube. The camera was set on a Z-mobile stage and affixed with two LED clamp lights. Depending on the desired image distance, a 20 mm long extension ring was also attached in front of the lens. The camera was situated about 20 cm from custom-built locomotion chambers that had transparent floors situated over an array of 90° reflecting prisms, allowing for simultaneous recording of lateral and ventral views. For calibration, the back walls and roofs of these chambers were lined by 5 mm^2^ paper. The program Photron Fastcam Viewer 4 was used for live transmission and recording. The frame rate was set to 250 fps and shutter speed to 1/500 s for most recordings; to account for rapid motion, however, these variables were set to 500 fps and 1/1000 s.

### 2.4. Data collection and analysis

Each kinematic recording consisted of a single, uninterrupted run that was as straight as possible, distant from the chamber walls, and centered over the prism array, thus allowing all tarsi to be seen in both views. Lighting and contrast were corrected via post-processing so that the tarsal outlines were clear. After multiple measurements were taken from a single individual, the specimen was assigned a number (i#) and immediately weighed using a calibrated analytical balance (ABS 80-4, Kern & Sohn, Germany), with the mass recorded in mg; deviations of up to 0.3 mg were found. Voucher specimens were preserved in 70% ethanol and retained in the Boudinot research collection (BEBC). The freeware program Tracker (Brown *et al*. 2023), which uses the open-source Physics Java Library (OSP), was used to calibrate each recording and to manually mark eleven different point masses frame-by-frame (Table 1). Point masses were tracked up to 320 consecutive images such that each measurement enabled the investigation of three stepping cycles (*e.g.*, Wahl *et al.,* 2015). To account for size scaling, several variables set to body length, which was measured linearly from the anterior apex of the head to the posterior apex of the metasoma in ventral view (Fig. S13).

**Table 1.** Overview of the tracked point masses.

| No. | Designation | Positioning explanation |
| --- | --- | --- |
| 1 | Head | Anteromedial point on head |
| 2 | Abdomen | Posteriormost point of abdomen |
| 3 | TarsusL1 | Left propretarsus |
| 4 | TarsusL2 | "" mesopretarsus |
| 5 | TarsusL3 | "" metapretarsus |
| 6 | TarsusR1 | Right propretarsus |
| 7 | TarsusR2 | "" mesopretarsus |
| 8 | TarsusR3 | "" metapretarsus |
| 9 | CoxaL1 (CoxaR1) | Left (or right) procoxa |
| 10 | CoxaL2 (CoxaR2) | "" mesocoxa |
| 11 | CoxaL3 (CoxaR3) | "" metacoxa |

**Table 2.** Taxon sampling overview. . Body size, basic performance, and discrete behavioral variables included. Abd. drag = obvious abdominal contact with floor made during recording. Knee posture = femorotibial joint raised above mesosomal dorsum in lateral view. Tar. slip = tarsal slippage observed. Discrete tokens: 1 = true, 0 = false. Average TCS values for genera with multiple samples indicated in parentheses.

| Taxon | ID | Mass (mg) | BL (mm) | Speed (mm/s) | DF (%) | TCS | Abd. drag | Knee posture | Tar. slip |
| --- | --- | --- | --- | --- | --- | --- | --- | --- | --- |
| **Hymenoptera: symphytan grade** |  |  |  |  |  |  |  |  |  |
| Superfamily Tenthredinoidea, Families Argidae, Tenthredinidae |  |  |  |  |  |  |  |  |  |
| *Arge* sp. | 12 | 24.1 | 11.1 | 14.3 | 82 | 0.00 | 1 | 0 | 1 |
| *Tenthredo* sp. | 11 | 30.6 | 12.2 | 22.8 | 80 | 0.00 | 0 | 0 | 0 |
| **Apocrita: parasitican grade** |  |  |  |  |  |  |  |  |  |
| Superfamily Ichneumonoidea, Families Braconidae, Ichneumonidae |  |  |  |  |  |  |  |  |  |
| *Bracon* sp. | 22 | 2.7 | 5.2 | 15.4 | 73 | 0.65 | 0 | 0 | 0 |
| (Ichneumonidae) | 18 | 6.6 | 9.0 | 24.6 | 77 | 0.03 | 0 | 1 | 1 |
| Superfamily Cynipoidea, Family Cynipidae |  |  |  |  |  |  |  |  |  |
| (Cynipidae) | 17 | 3.7 | 5.5 | 13.3 | 71 | 0.13 | 0 | 0 | 0 |
| Superfamily Evanioidea, Family Gasteruptiidae |  |  |  |  |  |  |  |  |  |
| *Gasteruption* sp. | 15 | 6.5 | 12.1 | 23.6 | 68 | 0.10 | 1 | 0 | 1 |
| **Apocrita: Aculeata: non-ants** |  |  |  |  |  |  |  |  |  |
| Superfamily Chrysidoidea, Family Chrysididae |  |  |  |  |  |  |  |  |  |
| *Chrysis* sp. | 7 | 8.5 | 7.6 | 22.0 | 75 | 0.13 | 0 | 0 | 0 |
| Superfamily Vespoidea, Family Vespidae |  |  |  |  |  |  |  |  |  |
| *Polistes* sp. | 23 | 62.9 | 16.7 | 55.3 | 72 | 0.14 | 1 | 0 | 0 |
| *Vespula germanica* | 4 | 74.7 | 15.3 | 55.8 | 65 | 0.26 | 0 | 0 | 0 |
| *Vespa crabro* | 6 | 328.4 | 26.2 | 78.5 | 69 | 0.01 | 0 | 0 | 1 |
| Superfamily Tiphioidea, Family Tiphidae |  |  |  |  |  |  |  |  |  |
| *Tiphia* sp. | 13 | 24.2 | 12.3 | 42.4 | 66 | 0.66 | 0 | 0 | 1 |
| Superfamily Apoidea, Families Crabronidae, Philanthidae, Halictidae, Andrenidae, Apidae |  |  |  |  |  |  |  |  |  |
| *Crabro* sp. | 19 | 33.7 | 12.5 | 41.6 | 65 | 0.42 | 0 | 0 | 0 |
| *Cerceris* sp. | 14 | 42.2 | 13.5 | 62.2 | 63 | (0.65) | 1 | 0 | 1 |
| *Cerceris* sp. | 20 | 8.3 | 7.6 | 31.4 | 63 |  | 1 | 0 | 1 |
| (Halictidae) | 8 | 4.8 | 6.9 | 27.0 | 62 | 0.47 | 0 | 0 | 0 |
| *Andrena* sp. | 5 | 73.5 | 16.2 | 37.8 | 71 | 0.50 | 1 | 0 | 1 |
| *Apis mellifera* | 3 | 74.3 | 18.2 | 61.3 | 69 | 0.31 | 1 | 0 | 1 |
| *Bombus terrestris* | 9 | 189.8 | 19.3 | 40.5 | 72 | 0.19 | 1 | 0 | 1 |
| **Apocrita: Aculeata: ants** |  |  |  |  |  |  |  |  |  |
| Superfamily Formicoidea, Family Formicidae |  |  |  |  |  |  |  |  |  |
| *Camponotus* sp. (w) | 42 | 12.3 | 10.3 | 30.8 | 79 | 0.24 | 0 | 1 | 0 |
| *Formica* cf. *fusca* (w) | 26 | 3.4 | 6.0 | 61.4 | 60 | 0.57 | 0 | 1 | 0 |
| *Lasius fuliginosus* (w) | 21 | 2.4 | 5.0 | 26.8 | 58 | (0.76) | 0 | 1 | 0 |
| *Lasius* sp. (w) | 36 | 2.2 | 4.3 | 79.2 | 58 |  | 0 | 1 | 0 |
| *Lasius* sp. (w-p) | 35 | 5.6 | 5.7 | 20.0 | 58 |  | 0 | 1 | 0 |
| *Lasius brunneus* (w) | 37 | 1.4 | 3.6 | 70.2 | 58 |  | 0 | 1 | 0 |
| *Lasius niger* (q) | 27 | 26.1 | 10.7 | 76.9 | 58 |  | 0 | 1 | 0 |
| *Lasius niger* (w) | 28 | 1.6 | 4.4 | 67.2 | 58 |  | 0 | 1 | 0 |
| *Myrmica* sp. (w) | 29 | - | 4.7 | 20.5 | 61 | (0.63) | 0 | 1 | 0 |
| *Myrmica* sp. (w) | 30 | 3.5 | 4.8 | 40.7 | 71 |  | 0 | 1 | 0 |

For the analysis of temporal stepping pattern, each recording was examined in similar to Humeau *et al.,* (2019): Tarsal movement was observed frame-by-frame and the image numbers for liftoff and touchdown were recorded. At least two complete contact phases with one swing phase included were tracked. Since some Hymenoptera dragged their hind legs over the ground, attention was paid to the onset of an active movement in the metatarsi; as soon as the tarsus left its original contact point, the swing phase was considered to start (lift-off time of Reinhardt & Blickhan, 2014, Wahl *et al.,* 2015). After recording liftoff and touchdown, the readings were converted to seconds depending on their frame rate and the durations of the swing and contact phases were plotted using stacked bar charts in white and black, respectively (*e.g.*, Zollikofer, 1994a; Grabowska *et al.,* 2012). Step patterns were subsequently scaled to one second. The duty factor was calculated from the relationship between swing time and contact time (Weihmann *et al.,* 2017; Wahl *et al.,* 2015):

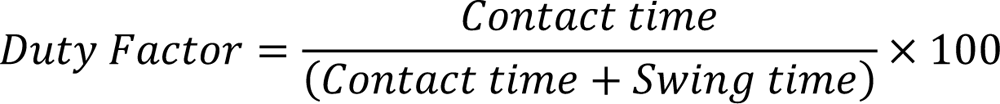

For the analysis of spatial stepping pattern, Tracker was used to quantify the triangles defined by the tarsal point masses when three tarsi (the middle and contralateral front and rear) were in the contact phase and the other three were in the swing phase (Zhao *et al.,* 2018). The times were determined in three consecutive steps, so that the tripodal leg groups alternated (Zollikofer, 1994a; Seidl & Wehner, 2008; Wahl *et al.,* 2015; Pfeffer et al., 2019) these steps were determined to begin when the standing middle leg was overtaken by the swinging isosegmental leg. The coordinates of the relevant tarsi (L1–R2–L3 or R1–L2–R3) from the respective three individual images were connected to form triangles. The point masses for the head and abdomen were plotted to serve as a body-fixed coordinate system and the lengths of the individual lines and the distances between the point masses were calculated, allowing for the determination of body length, stride length, and step width. The latter indicates the lateral distance from the center of the body to the touchdown point of the tarsus in mm and comprises the mean values each isosegmental pair of legs (L1–R1, L2–R2, L3–R3). The distance covered by the central point between the head and abdomen along the longitudinal body axis was used to calculate stride length in the time span from the touchdown of the middle tarsus to the next touchdown of the same tarsus (Pfeffer *et al.,* 2019; Wahl *et al.,* 2015; Zollikofer, 1994a).

Tripod coordination strength (*e.g.*, Wosnitza *et al.,* 2013) was determined using the following formula:

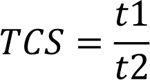

Here, t1 designates the period in which all three legs of a tripod are in the swing phase and t2 the period in which at least one of these three legs is in a standing phase, thus t1 ≤ t2 always applies. A TCS value of 1 would, therefore, mean that the tripod coordination is perfectly synchronous. The closer the TCS value approaches 0, the less synchronous the swing phases of the tripod are (Pfeffer *et al.,* 2019; Merienne *et al.,* 2020). TCS was calculated for both tripod pairs (L1–R2–L3, R1–L2–R3) over a three-step interval. The mean value was then formed from all steps of both tripod pairs.

For range of motion analyses, each leg axis was defined between the coxa and tarsus. It was assumed for simplicity that the insect leg moves one plane (“leg plane”; *e.g.*, Cruse & Bartling, 1995; Seidl & Wehner, 2008); out-of-plane bending at the trochanterofemoral joint was not accounted for (see OpenSim model in Arroyave-Tobon *et al*., 2022). The angle between the longitudinal body axis (BLA) and the leg axis at the anterior (AEP) or posterior (PEP) extreme positions was calculated in ventral view (Fig. S13). The minimum leg angle is between BLA and AEP and the maximum between BLA and PEP. Due to variation in leg movement within a run, the range of motion was determined over at least two steps. The resulting tarsal trajectory (*Schneeengel*) plots were scaled to the body length of the individual and mirrored if necessary. To quantify differences between ant and non-ant samples, estimation graphics (Ho *et al*., 2019) with median differences (Δ), 5000 bootstrap resampled distributions, and bias-corrected 95% confidence intervals (CIs) were used in R 4.0.2 (R Foundation for Statistical Computing, Vienna, Austria) via the package dabestr v0.3.0.

## 3. Results

### 3.1. Temporal stepping pattern

Across the sampled Hymenoptera both the average and the spread of measured speeds was significantly higher for ants than non-ants (Fig. 1A). While the non-ants had an average relative speed of 2.9 ± 2.03 BL/s (body length per second), the ants achieved 9.6 ± 5.85 BL/s. The fastest individual was a worker *Lasius* (i37) at 19.6 BL/s, which is ≥ 10 x as fast as the sampled sawflies (*Arge*, 1.3 BL/s; *Tenthredo*, 1.9 BL/s). The highest absolute speed was achieved by the sampled Vespidae (63.2 mm/s) with a maximum of 78.5 mm/s, while the Cynipidae had the lowest value (13.3 mm/s). As average walking speed is the product of stride length and step frequency, it is notable that some genera such as *Arge* or *Gasteruption* took very short steps while others, the sampled Formicidae in particular, moved with relatively large steps. Ants achieved an average of 0.70 ± 0.17 SL/BL (stride length by body length), a significantly higher value than non-ants with 0.51 ± 0.09 SL/BL (Fig. 1A). The maximum stride length in relation to body length was also measured in *Lasius*, which at 96% corresponded to almost the entire body length. The exception among sampled *Lasius* was the parasite infested individual (i35), which was the slowest of all sampled Formicidae (20 mm/s) and the lowest step frequency among sampled congeners (8 Hz). The ants, otherwise, achieved the highest average step frequency of all sampled Hymenoptera (Figs. 2, S15, S16), with a value of 10.3 Hz (*i.e.*, 10.3 steps per second), corresponding to an increase of 222% over the lowest values, which were observed for the sampled Tenthredinoidea (3.2 Hz).

**Fig. 1.**
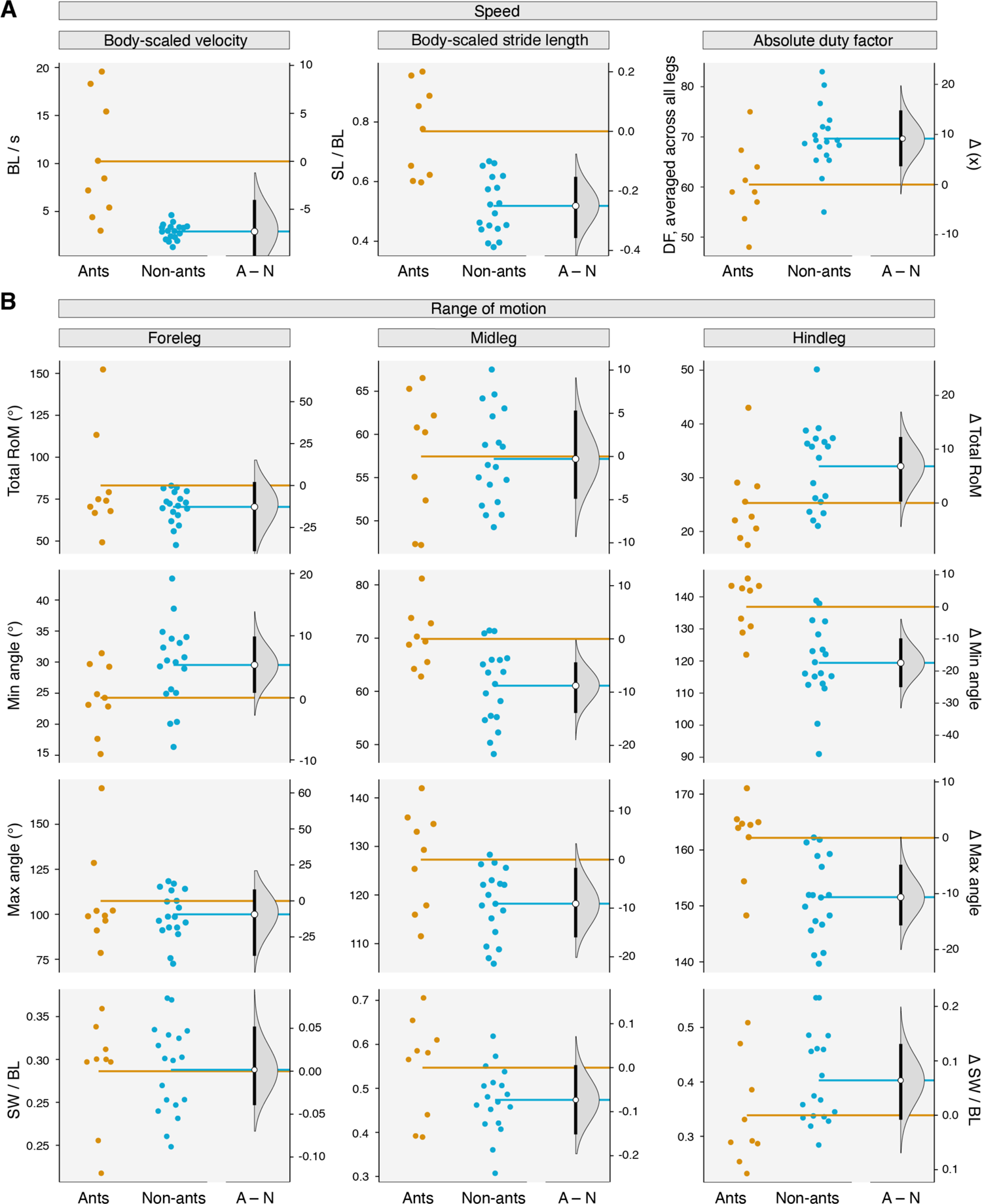
Temporal and spatial comparisons of ant and non-ant gaits. (A) Relevant speed metrics, organized by measurement. (B) Relevant range of motion metrics, organized by leg then my metric. Sample sizes based on count of recorded individuals: Ants n = 9 (parasitized specimen, i36, excluded); non-ants n = 18. In both A and B, grey distributions represent bootstrapped ranges from the non-ant sample, black bars represent 95% CIs, and open circles indicate non-ant means. For comparisons in which the 95% CI does not overlap with zero, the result can be interpreted as a significant difference at an alpha of 0.05. Plots not to the same scale. Note that body mass has a range of 1.6–26.1 mg for ants and 2.7–328.4 mg for non-ants.

**Fig. 2.**
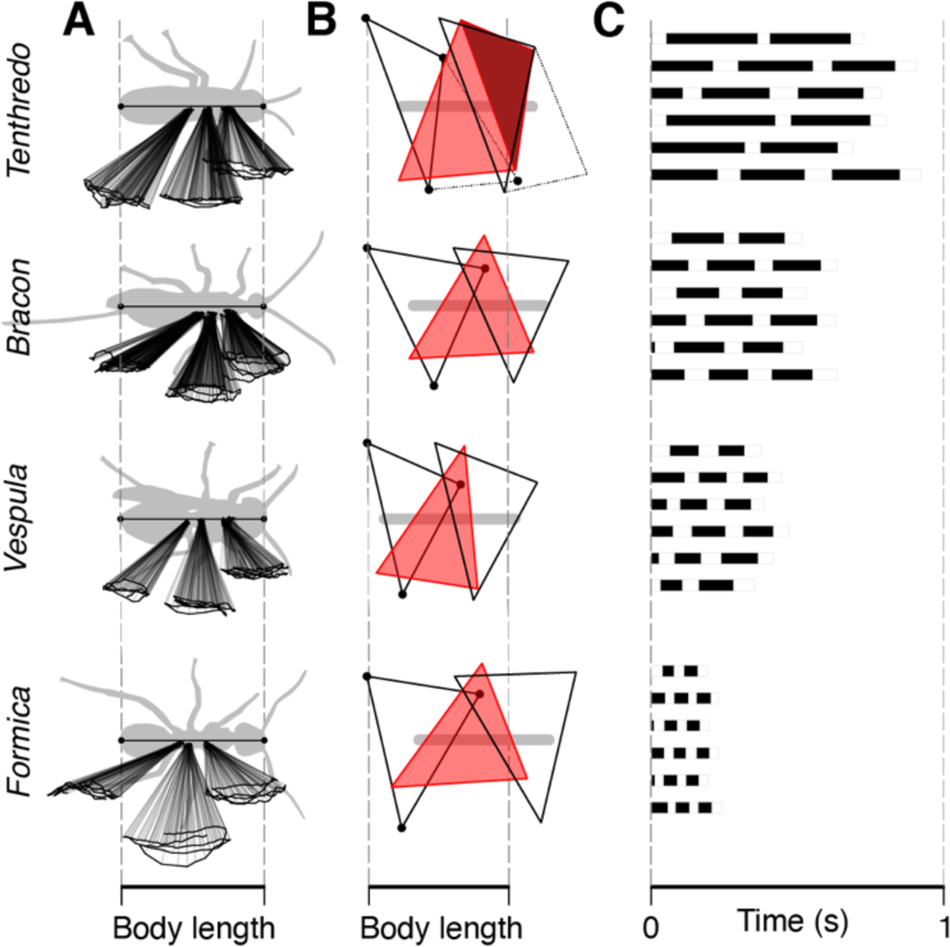
Representative locomotory patterns of Hymenoptera scaled to body length or time. (A) Leg-by-leg range of motion. (B) Spatial gait pattern; *Tenthredo* plotted with additional triangles due to the observed tetrapedalism. (C) Temporal gait pattern averaged across all measurements per individual. The four taxa were chosen to represent, from top to bottom: “symphyta”, “parasitica”, non-ant Aculeata, and Formicidae. For visualization of results across all taxa, see Fig. S16.

Across most sampled superfamilies, the swing time was between 0.05 and 0.06 seconds (Figs. 3, S15, S18); only for Chrysidoidea and Formicoidea was this value on average 0.04 seconds. Values for all legs were similar among all sampled taxa, with the exception of the forelegs of *Camponotus* and the hindlegs of *Tenthredo*, *Bracon*, *Cerceris*, and *Bombus*, which had a relatively long swing phase. In contrast, variation in contact time was much more striking (Figs. 3, S18). The average contact time of Formicoidea was 0.07 seconds, which was the lowest across all sampled Hymenoptera and was 72% less than that of Tenthredinoidea (0.25 s). The Tenthredinoidea also walked with the highest duty factor—the contact phase here comprised 81% of the entire gait cycle on average, while the Formicoidea lowest average duty factor of all sampled Hymenoptera at 63%. Ants had significantly lower duty factors overall (Fig. 1A). The shortest swing (0.02 s) and contact times (0.03 s) as well as the lowest duty factor (54%) among all samples was observed for *Lasius*, while *Camponotus* stood out among the sampled ants as it had the longest contact phase by far and the highest duty factor (79%) (Tables S6–8).

**Fig. 3.**
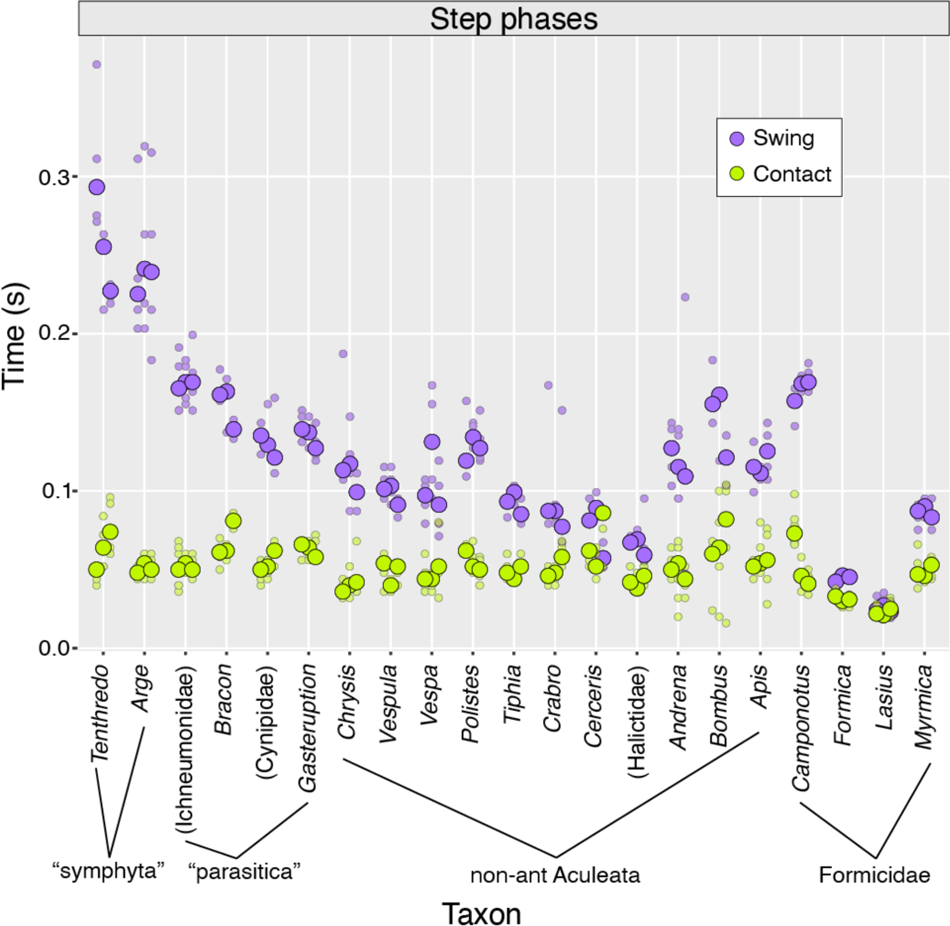
Direct comparison of swing versus contact times across sampled Hymenoptera, aggregated by genus. Median values for each leg represented by large dots, each individual measurement by smaller dots. See Fig. S18 for leg-by-leg comparison.

### 3.2. Spatial stepping pattern

The temporal stepping pattern also reflects the gait used to move. All individuals with the exception of the *Tenthredo* and *Arge* showed features of the alternate tripod gait (Figs. 2, S15, S16; Tables S6, S7). The tripod coordination strength (TCS) of these sawflies was invariably 0 because at no point in time of the evaluated steps were three legs in a swing phase, thus four legs always remained in the contact phase, resulting in a tetrapod gait. Among other taxa, the gait varied without this extreme. The TCS values for *Vespa* (0.01) and Ichneumonidae (0.03) also recorded a conspicuously low value, running in a mainly asynchronous tripod gait. Notably, the locomotion of the sampled *Vespa* was interrupted by wing flapping behaviour in all measuring periods. Specimens of the genera *Lasius* (0.76), *Tiphia* (0.66), *Cerceris* (0.65), *Bracon* (0.65), and *Myrmica* (0.63), in contrast showed a very high tripod coordination.

Overall, the range of motion of all superfamilies was greatest for the forelegs, with an average angle of about 72°, followed by the mid- (57°) and hindlegs (29°). The smallest average touchdown or minimum angle for the foreleg was 20° (Chrysididae), 56° (Apoidea) and 58° (Gasteruptiidae) for the midleg, and 113° (Tiphiidae), 114° (Apoidea), and 116° (Tenthredinoidea) for the hindleg. The largest such values for the fore-, mid-, and hindlegs were observed for Tenthredinoidea (41°), Cynipoidea (71°), and Formicoidea (138°), respectively. For tarsal lift-off, the smallest average maximum angle values for the forelegs were 89° (Vespidae) and 93° (Chrysididae); they were 109° for the midlegs (Gasteruptiidae) and 147° for the hindlegs (both Tenthredinoidea and Tiphiidae). The largest such angles were 118° for the forelegs (Tenthredinoidea), 127° for the midlegs (Formicoidea and Ichneumonoidea), and 163° for the hindlegs (Formicoidea).

Comparatively, the ants were very similar to other sampled Hymenoptera with respect to foreleg and hindleg total range of motion as well as foreleg maximal angle (Figs. 2, S15, S16). They differed significantly, however, in having a smaller average foreleg minimum angle, in all measures of midleg range of motion, and in having more extreme minimal and maximal hindleg angles (Figs. 1B, 2, S15, S16). As for step width, the sampled ants were indistinguishable for forelegs, but had on average wider midleg and narrower hindleg locomotory stances (Figs. 1B, 2, S15, S16). The greatest step width relative to body length for the forelegs (37%) and hindlegs (49%) was recorded from Tenthredinoidea, while the lowest relative step width values overall were recorded for the gasteruptiid (20% foreleg, 31% midleg, 30% hindleg).

## 4. Discussion

To date, no publication has examined locomotion across the Hymenoptera, let alone in a phylogenetic context. More broadly, investigations of hymenopteran gait or stepping patterns are seldom, with the only non-ant being *Apis mellifera*, and usually only concern a single species, with a few exceptions that focus on the walking or running of closely related ants. These studies have taken the locomotory capacity of ants as granted, as our results show for the first time that this is unequivocally derived (Fig. 4). Moreover, we find—to our surprise—that the wide-waisted wasps (“symphyta”) in our sampling have a consistent tetrapod gait. Below we discuss our results based on our expectations, as outlined in the introduction: (4.1) the tripod gait will be dominant, (4.2) other Hymenoptera will be able to run, and (4.3) ants will have modified ranges of motion. Additionally, we address two unanticipated considerations arising from our observations, namely the sequence of leg liftoff and attachment variability across our sample (4.4).

**Fig. 4.**
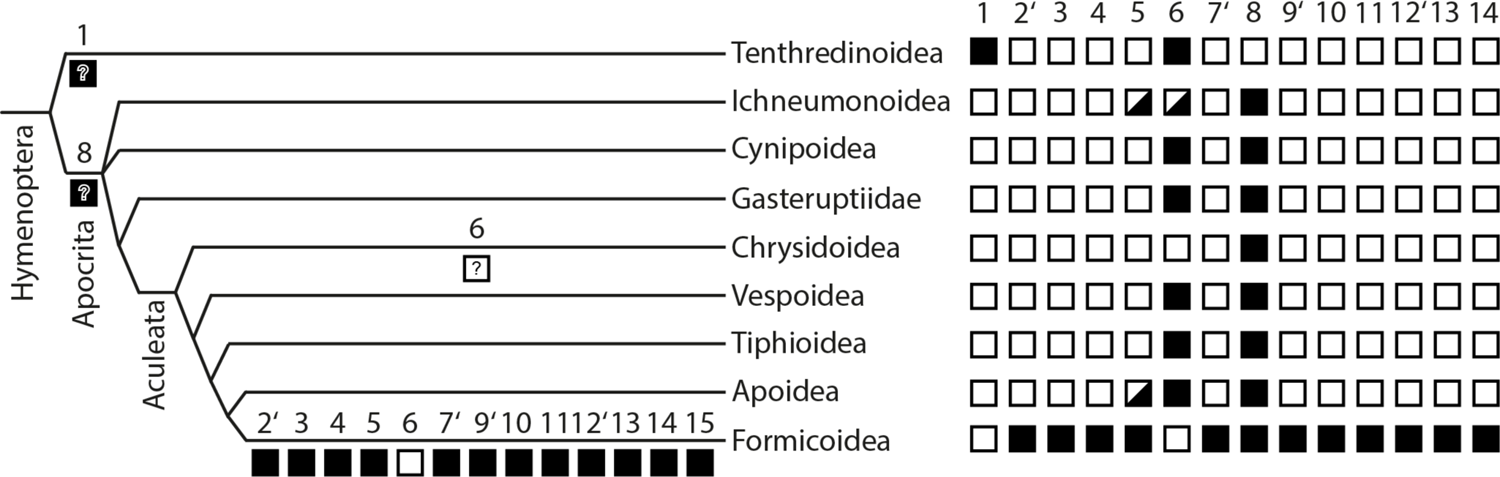
Phylogenetic/evolutionary summary of key results. Parsimonious interpretations of functional and structural characters for the sampled Hymenoptera indicated at the respective nodes, emphasizing distinctions of Formicidae. Apostrophes by character numbers indicate previously determined derivations of Formicidae. Characters (open-box conditions first, closed-box second): **(1)** Dominant gait observed: tri- vs. tetrapod; **(2’)** wing dimorphism/polyphenism: absent vs. present; **(3)** locomotion: perambulatory (< 5 BL/s, < 0.7 SL/BL) vs. cursorial (≥ 5 BL/s, ≥ 0.7 SL/BL); **(4)** observed duty factor: high (≥ 60%) vs. low (< 60%); **(5)** femorotibial joints raised above mesosomal dorsum during locomotion: not observed vs. observed (*note*: among non-ants, observed in an ichneumonid and the apoid genus *Cerceris*); **(6)** abdominal apex dragging: not observed vs. observed (*note*: trail laying for ants excluded); **(7’)** procoxotrochanteral articulation: dicondylic, open vs. monocondylic, closed; **(8)** foreleg relative step width: broad (> 0.4 SW/BL) vs. narrow (≤ 0.4 TW/BL); **(9’)** proximal mesocoxal articulation: wide vs. narrowed, ball-in-socket-like; **(10)** relative midleg range of motion: anterior (< 73° AEPm, < 130 PEPm) vs. posterior (≥ 73° AEPm, ≥ 130 PEPm); **(11)** maximum midleg step width: narrow (< 0.6 SWm) vs. broad (≥ 0.6 SWm); **(12’)** proximal metacoxal articulation: wide vs. narrowed, ball-in-socket-like; **(13)** relative hindleg range of motion: anterior (< 140 AEPh, < 162 PEPh) vs. posterior (≥ 140 AEPh, ≥ 162 PEPh); and **(14)** maximum hindleg step width: broad (≥ 0.4 SWh) vs. narrow (< 0.4 TWh). For Char. 1 see Fig. S16, for Chars. 5, 6 see Table 2, for Chars. 7, 9, 12 see Boudinot (2015) and Boudinot *et al*. (2022), and for Chars. 10, 11, 13, and 14 see Fig. 1 and Table S7. Further sampling of “symphyta” is necessary to resolve hymenopteran groundplan patterns. Ant autapomorphies may be found to be synapomorphies with explicit sampling of taxa with wingless morphs.

### 4.1. Is the tripod gait universally dominant among Hymenoptera?

Not necessarily. To our surprise and in contrast to all other sampled Hymenoptera, none of the sampled symphyta (n = 2 individuals; Argidae, Tenthredinidae) walked with an alternating tripod gait (Figs. 2, S15, S16); the TCS values of both taxa were 0. Future comparative study with expanded sampling is an absolute necessity. Regardless, across all seven measurements, no tripod gait could be identified, since at least four legs were always on the ground, as expected for a tetrapod gait (Grabowska *et al.,* 2012; Chen *et al.,* 2012). The movement of *Arge* and *Tenthredo* resembled a “metachronal wave gait” because it started in the hind legs and transferred posterior to anterior from the hind- to the forelegs (Ambe *et al.,* 2018); their gait was unusual as the swing phases of the front legs were sometimes simultaneous, such that the tetrapod gait was more irregular (Grabowska *et al.,* 2012). It is possible that the sampled symphyta simply had particularly low speeds, as insects are known to change their gait depending on speed (Merienne *et al.,* 2020) and that a tetrapodal rather than tripodal gait tends to be taken during walk (Grabowska *et al.,* 2012; Ambe *et al.,* 2018; Arroyave-Tobón *et al.,* 2022). However, it is possible that other dynamics are at play, such as skeletomuscular modification. The Hymenoptera are defined, in part, on modifications of their pleurosternal thoracic regions (*e.g.*, Gibson 1993; Vilhelmsen 2000a), which have unknown functional consequences. Without data on the further six superfamilies of “symphyta”, it is impossible to determine the character polarity of tetrapodal locomotion at the base of the hymenopteran phylogeny. *I.e.*, tetrapodal locomotion may have evolved in the ancestor of the Hymenoptera with secondary gain of tripodal locomotion in the ancestor of the Apocrita, or a four-legged stance phase may define Tenthredinoidea or be derived therein (Fig. 4).

### 4.2. Can other Hymenoptera run?

Although we expect that other Hymenoptera have the capacity to run (*e.g.*, Schmidt & Blum 1977), we found that cursoriality is neither widespread nor likely an ancestral condition of the order (Fig. 4). Specifically, all sampled non-ant Hymenoptera clustered tightly at an apparent walking pace with respect to locomotory speed as scaled to body length, whereas this measure shows that ants were clearly able to change their speeds from a walk to a run (Fig. 1A). Correlated with the greater relative speeds achieved by ants are their posture, with their femorotibial joints elevated above their thoracic dorsum (Fig. 4: Char. 5), and their elevated metasomata, which did not contact the walking surface, excepting trail-laying behavior (Fig. 4: Char. 6). These results indicate that adaptation from flight dependency to surface locomotion dominance in the ancestor of the ants—the sky-to-ground transition—had complex anatomical, functional, and performance consequences.

While our total sample size may be limited (n = 28 individuals), the phylogenetic breadth of our sampling spans the topological backbone of the order and > 250 million years of evolution (*e.g.*, Peters *et al*. 2017; Boudinot *et al*. 2022), strongly indicating that the locomotory capacity of ants is derived. It is possible that some of the non-ant Hymenoptera achieved a run with respect to mechanical energy and that this was masked by the gait, which may have been a grounded run, Groucho run, or compliant walking, as observed for *Formica polyctena* (Reinhardt & Blickhan 2014). Regardless, the robustness of our observations in terms of body-scaled speed and stride length plus absolute duty factor is enhanced by the coincidental sampling of a queen *Lasius* and a parasite-infested congeneric worker (Table S8, Fig. S19), as the former achieved similarly high speeds to healthy workers despite the massive mesosoma, the large abdomen, and overall longer body. In contrast, the infested worker displayed very short stride lengths, which may be a symptom of disease (Hölldobler, 1952; Kraus & Page, 1995; Klimov *et al.,* 2007).

More generally, in order to increase walking speed, animals must increase their stride length and/or reduce the duty factor (Raibert *et al*. 1989; Alexander, 2003; Nishii, 2006; Wittlinger et al., 2007). Across all sampled Hymenoptera, the average duty factor was very high (70%) and was highest in the Tenthredinoidea (81%). Compellingly, the lowest duty factor was found in ants, which averaged about 63% (Fig. 1A). In other studies, too, the duty factor of ants was similar; for example, *Colobopsis schmitzi* (formerly *Camponotus*) had an average duty factor of 60% (Bohn, 2007; Bohn *et al.,* 2012), *Formica polyctena* achieved a duty factor of 65% with a speed of 80 mm/s (Reinhardt & Blickhan, 2014), and *Aphaenogaster subterranea* were also measured as having a duty factor of 66% on a straight, smooth surface (Humeau *et al.,* 2019). The duty factors were still high enough to enable a stable tripod gait at fairly high speeds (Nishii, 2006; Ting *et al.,* 1994; Seidl & Wehner, 2008), however, as ants had by far the longest stride length in relation to body length as well as the highest speed of all sampled taxa.

With respect to stride length, the genus *Lasius* was characterized by particularly long strides among our sampled Formicidae. One individual in particular (*L. brunneus*, i37) had the smallest body length yet achieved the highest relative speed, with a stride that was 96% of its body length (Fig. S19), similar to the *Cataglyphis fortis* sampled by Wahl *et al.,* (2015) running at a speed of 95.2 mm/s, albeit far less than that species maximum speed (596 m/s, 60 BL/s). Notably, the non-ants that had the longest stride lengths (*Cerceris* i14, *Polistes*, *Bracon*) raised their hindleg femorotibial joint above the dorsal abdominal margin in lateral view, which otherwise only occurred in the sampled ichneumonid and the ants. Quantification of lateral view kinematics, interleg allometry and stride length would be a reasonable next step to determine the degree of correlation between leg and stride length across Hymenoptera (*e.g.*, Reinhardt & Blickhan, 2014; Wittlinger *et al.,* 2007).

We further observed that changes in step frequency across our dataset were mainly due to changing contact times, which varied between 0.07 and 0.25 seconds, while swing times varied only between 0.04 and 0.06 seconds (Fig. S18, Table S7). Swing times were thus relatively stable as generally observed (Nishii, 2006), supporting the assumption that Hymenoptera, like other insects, have similar central pattern generators and motor neurons that trigger muscle activity in the legs and also coordinate the tripod gait (Kukillaya *et al.,* 2009; Guo *et al.,* 2013; Ambe *et al.,* 2018; Barrio *et al.,* 2020). Ants were, however, observed to reduce the duration of their swing phase in order to achieve maximum speed (Figs. S18, S19), as also recorded for *Formica rufa* (Reinhardt & Blickhan, 2014).

### 4.3. Do ants have a distinct range of motion?

Our results strongly indicate that, as expected based on comparative morphology (Boudinot 2015), Formicidae have distinct and evolutionarily derived ranges of motion during locomotion. While several families had more extreme measurements for certain variables, the significant differences in leg mobility of ants relative to other Hymenoptera (Fig. 1B) are meaningfully interpreted as consequences (or at least correlates) of their structural distinctness. Notably, we did not see major differences for the forelegs despite their autapomorphic transition from coxotrochanteral dicondyly to monocondyly in the ancestor of ants (Boudinot 2015; Boudinot *et al*. 2022). Rather, ant forelegs performed as expected for Apocrita, for which clade the range of motion itself may be derived, as the sampled sawflies had the broadest foreleg step widths and were unable to draw these legs close to the midline of the body (Fig. 4: Char. 1). Mid- and hindleg performance, in contrast, is where ants were clearly distinct, achieving the overall widest midleg and narrowest hindleg step widths as well as the furthest posterad/mediad shifts of the minimum and maximum angles for both pairs of legs. Broad midleg track width in relation to the other legs in ants has also been observed by Merienne *et al.,* (2020). Given the important roles of the mid- and hindlegs across insects (Full & Tu, 1991; Cruse, 1976; Reinhardt & Blickhan, 2014; see also Wöhrl *et al.,* 2017, Merienne *et al.,* 2020) it is possible that these relative step widths maximize propulsion via the hindleg and, for the midleg, increase the torque generated with respect to the normal (yaw) axis for quick direction changes. In other words, a wide midleg stance may increase the length of the lever arm. Optimization of torque may represent the central mechanical explanation for the extreme and derived ball-in-socket coxothoracic articulation of the meso- and metathoracic segments of ants (Boudinot 2015) via conservation of angular momentum and shortening of the inlever relative to the outlever. Form-function modeling will address this hypothesis and may reveal additional axes of anatomical and locomotory variation.

### 4.4. Considerations arising

Based on our observations, two unanticipated considerations arise: Are midlegs first to lift and hindlegs last, and is attachment variable across Hymenoptera?

#### Lifting sequence

Merienne *et al.,* (2020) observed in the harvester ant *Messor barbarus* that it tended to perform its tripod swing phases in a specific order. The lifting of the legs usually happened in the following sequence: midleg → foreleg → hindleg. Occasionally the mid- and forelegs swapped with each other, but the hindleg was always moved last. Most of the ant genera considered here conformed to the hindleg-last expectation, although the foreleg → midleg → hindleg order was the most common (n = 12). Among our sample, only the non-ant species of *Cerceris* and *Bracon* were observed to begin with a hindleg swing phase; these genera also had a pronounced tripod coordination of 0.65 TCS. Otherwise, it also happened that the hind leg was moved second (*Lasius*, *Bombus*, *Chrysis*) or that no distinct order emerged (*Apis*, *Andrena*). The study of *Camponotus fellah* by Tross *et al.,* (2022) showed that the order can change depending on the speed; only above a certain speed did the *Camponotus* move its hind leg first in the tripod. This may be related to the propulsion unit of the legs (Reinhardt & Blickhan, 2014; Merienne et al., 2020), as the *Camponotus* might have needed more push off from the hind legs to gain more speed, which in turn affected the tripod order. Regardless, our results indicate that lifting sequence is variable among Hymenoptera and requires further sampling to explain.

#### Attachment variability

It is known that friction and adhesive tarsal attachment structures vary across Hymenoptera (Schulmeister, 2003; Beutel *et al*., 2020; Boudinot *et al*., 2021). Across all sampled taxa, the attachment properties of Formicidae were the most pronounced, as they ran quickly on all surfaces of the measuring chamber, a behavior not observed in other groups. Unlike bees and wasps, ants use a hydraulically driven unfolding mechanism of the arolium for this purpose (Beutel *et al.,* 2020; Federle *et al.,* 2001). In addition, it has been shown that ant aroliae develop a very high adhesive force and can also be unfolded passively without the action of the claw flexor muscle (Endlein & Federle, 2008, 2013), although ant aroliae differ in their properties such as size or surface structure with correlation to their natural history, such as reliance on climbing (Wöhrl *et al*., 2021; Billen *et al*., 2017). We also observed that other taxa had more limited attachment ability. Most of the sampled bees, for example (*Andrena*, *Apis*, and *Bombus*), slipped on the locomotion chamber floor despite their ability to grip petal surfaces (*e.g.*, Bräuer *et al.,* 2017), while the sampled vespids were clearly much more capable of attaching to various surfaces. The fact that some bees were scooting suggests that they were walking on their claws with the arolia folded in. Nevertheless, genera belonging to Apoidea had an average of 0.42 ± 0.1 TCS, thus showing a more temporally consistent tripod gait than Vespoidea, which had an average of 0.14 ± 0.1 TCS (Tables S5, S6).

## 5. Conclusion and outlook

Based on our performance measures, our results show for the first time that ants have evolutionarily specialized locomotory capacity. The evidence for ant cursoriality was reflected in numerous parameters, including their relatively long strides, higher step frequency and speed, lower swing time and duty factor, strong tripod coordination, and absence of slipping. These results were anticipated but not predicted by prior comparative morphological work. The evolutionarily derived ball-in-socket like coxothoracic joints of ant mid- and hindlegs appear to modify the range of motion for these legs, as these legs were observed to have broad and narrow step widths, respectively, and both leg pairs could extend further posterad than other Hymenoptera. We therefore hypothesize that these derived range of motion parameters are central to the sky-to-ground transition that characterizes ants as cursorial wasps with winged-wingless polyphenism. The exact mechanisms causing the distinct range of mid- and hindleg motion for ants remains to be explored, while the consequences of the uniquely derived monocondylic articulation of the ant procoxotrochanteral joint is as yet unexplained. Contrary to expectation, we observed that the sampled sawflies walked with a strict tetrapod gait, raising questions about the evolutionary dynamics of locomotion from the phytophagous “symphyta” to the carnivorous Apocrita. Furthermore, derivation of the apocritan foreleg coxotrochanteral joint is suggested by the narrow step widths of this segment for narrow-waisted wasps relative to sawflies.

Overall, our work establishes the Hymenoptera as a model system for studying the evolution of locomotion. The parameters and dynamics of the form-function relationship are expected to vary significantly across the order, given the broad ecological space occupied by adults (*e.g.*, Gauld & Bolton, 1988) and the exceptionally variable thoracic skeletomuscular anatomy of Hymenoptera relative to other insect clades at ordinal level (*e.g.*, Vilhelmsen, 2000b; Vilhelmsen *et al*., 2010; Friedrich & Beutel, 2010). Within Hymenoptera, the widespread derivation of sexual wing dimorphism and even within-sex diphenism (*e.g.*, Hanna & Abouheif 2021) provides substantial opportunity to study the evolution of locomotor systems across the transition from flight to walking dependency (*e.g.*, Reid 1941, Peeters *et al*., 2020), as well as to more unusual modes such as wood boring (*e.g.*, Khalife *et al*., 2018), saltation (Baroni Urbani *et al*., 1994; Ye *et al*., 2020), and swimming (*e.g.*, Yanoviak & Frederick, 2014; Schultheiss & Guénard, 2021). By linking phylogeny and mechanics through comparative kinematics, our results indicate that ants have achieved a distinct engineering solution to rapid running via high locomotory clearance and a knees-above-the-back posture, in contrast to the low and knees-forward architecture of cockroaches and beetles. Whether this solution is globally unique or simply a local optimum awaits future study. It is therefore certain that further surveys of locomotory biodiversity will yield fresh insight into the natural experiment of animal evolution.

## Acknowledgments

We thank Prof. Dr. Manuela Nowotny, Prof. Dr. Rolf G. Beutel, and PD Dr. Hans Pohl for the opportunity to work with the Animal Physiology and Entomology groups at Uni Jena. For helpful suggestions and constructive feedback on an earlier draft of this work, we thank Prof. Drs. Nowotny, Phil Ward, and Jack Longino, and Dr. Stefan Schöneich. For assistance in the field and for providing additional specimen identifications, we thank PD Dr. Pohl, Michael Weingardt, Di Li, and Daniel Tröger. An unpublished earlier draft of this work was completed in partial fulfillment of the requirements for *Wissenschaftliche Hausarbeit zur Ersten Staatsprüfung für das Lehramt am Gymnasium* for VR. BEB acknowledges support from the Alexander von Humboldt Stiftung via a Research Fellowship (2020–2022) for which Prof. Dr. Beutel and PD Dr. Pohl graciously sponsored his stay in Jena. BEB further acknowledges Prof. Dr. Dr. Martin Fischer and Prof. Dr. Andreas Hejnol for supporting his work at the Zoologisches Institut.

## Funding

– BEB was supported by a Research Fellowship from the Alexander von Humboldt Stiftung (2020–2022).

## Author contributions

– Conceptualization: **BEB**, **TW**.

– Methodology: **VR**, **BEB**, **TW**.

– Software: **TW**.

– Validation: **VR**, **TW**.

– Formal analysis: **VR**, **BEB**, **TW**.

– Investigation: **VR**, **BEB**, **TW**.

– Resources: **BEB**, **TW**.

– Data curation: **VR**, **BEB**, **TW**.

– Writing (original draft): **VR**, **BEB**.

– Writing (review, editing): **VR**, **BEB**, **TW**.

– Visualization: **VR**, **BEB**, **TW**.

– Supervision: **BEB**, **TW**.

– Project administration: **BEB**, **TW**.

– Funding acquisition: **BEB**.

## Competing interests

– The authors declare no known competing interests.

## Data availability

– Kinematic videos, output data, and R scripts will be made available at Zenodo after acceptance.

## Supplementary Information

### S1. Detailed methods

#### S1.1. Kinematic setup

As in Reinhardt & Blickhan (2014) and Wöhrl *et al.,* (2021), we used the high-speed Photron Fastcam SA3 camera, which had to be adjusted and calibrated beforehand. To dissipate heat the camera was placed crosswise on four aluminum framing extrusions. The extrusions were be fixed parallel to each other on a wooden panel, which in turn was located on a laboratory lifting platform (Fig. S1). Two LED clamp spots were attached to the wooden panel itself and their luminous flux of 220 lumen each was aligned to the measuring area. The camera lens from the manufacturer Schneider Kreuznach consisted of a C-mount lens adapter, a Makro-Unifoc 12 focusing tube and a Componon-S enlarging lens (Table S1). Depending on the desired image distance, a 20 mm long extension ring was also attached in front of the lens.

**Fig. S1.**
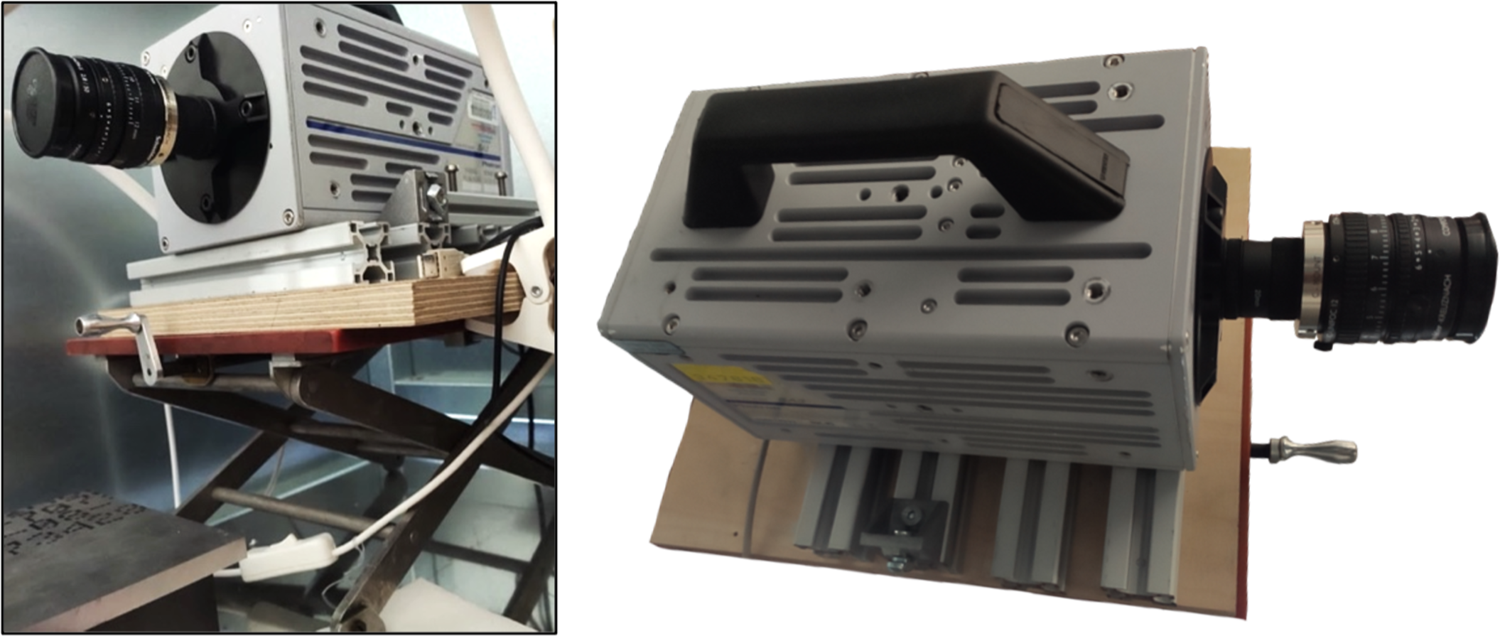
Frontal view of the high-speed camera on framing extrusion platform atop the laboratory jack (left) and the camera in the top view (right). Photos: Vincenz Regeler, August 10, 2022.

**Table S1.** Overview of the materials of the camera construction.

| Material or device designation | Manufacturer |
| --- | --- |
| Fastcam SA3 model 60K-M3 (high speed camera) | Photron; San Diego, CA, United States |
| 6.5mm lens adapter C mount | Jos. Schneider Optical Works GMLH; Bad Kreuznach, Germany |
| 17.4 – 29.4 mm focusing tube Makro-Unifoc 12 | Schneider-Kreuznach |
| 50mm f/2.8 Componon-S Enlargement Lens | Schneider-Kreuznach |
| 20 mm intermediate ring | Schneider-Kreuznach |
| "Nävlinge" clip-on LED spots | Inter IKEA Systems BV; Delft, Netherlands |
| Laboratory lifting platform (platform area 320 mm x 250 mm) |  |
| Aluminum profile (200 mm x 30 mm x 30 mm) |  |
| Wooden panel (300mm x 250mm x 20mm) |  |

The locomotion chamber was situated on a laboratory jack with an adjustment range of 60 to 250 mm at about 200 mm in front of the camera. The chamber consisted of a building board and plastic clamping blocks, which was used to fasten a row of three reflecting prisms. In addition, a transparent glass plate was lain horizontally on top of the chamber as a walking surface and observation panel (Fig. S2). The chosen prisms were right-angled isosceles triangles, and they were positioned such that the light could enter through the cathetus surfaces and were totally reflected and deflected by 90° on the hypotenuse surface. With this reflection function, the prisms served as an optical instrument for image generation instead of a mirror. As a result, in addition to the lateral view, the ventral view of the insect subjects could be recorded with a single camera, as in the study by Wosnitza *et al.,* (2013).

Once the test setup was prepared, the measuring range and camera could be adjusted to the desired working height using the jacks. For this purpose, in the live transmission of the camera program Photron Fastcam Viewer 4 (PFV 4), care was taken to ensure that the middle of the glass plate front was in the middle of the image, and that the top of the glass plate could still be seen, such that the tarsi in a lateral view are not obscured. The horizontal alignment of the measurement area and the camera could then be checked with a spirit level. Slight deviations could be helped with some modeling clay or by readjustment of the clamping blocks. To align the measurement area with the camera, the spirit level was also held against one side of the camera and at the same time against the side of the glass plate. A piece of corrugated cardboard about one centimeter in size was then placed on the glass plate above the prisms, *i.e*., on the “region of interest”, so that the camera could focus on it from two perspectives (directly from the front and indirectly through the prism below). The focusing tube of the camera could be adjusted in such a way that the sharpest possible image of the corrugated cardboard was obtained from both perspectives (Fig. S3). To minimize the effect of distance between the subject and the camera, the chamber was moved to the edge of the jack (Fig. S2).

**Fig. S2.**
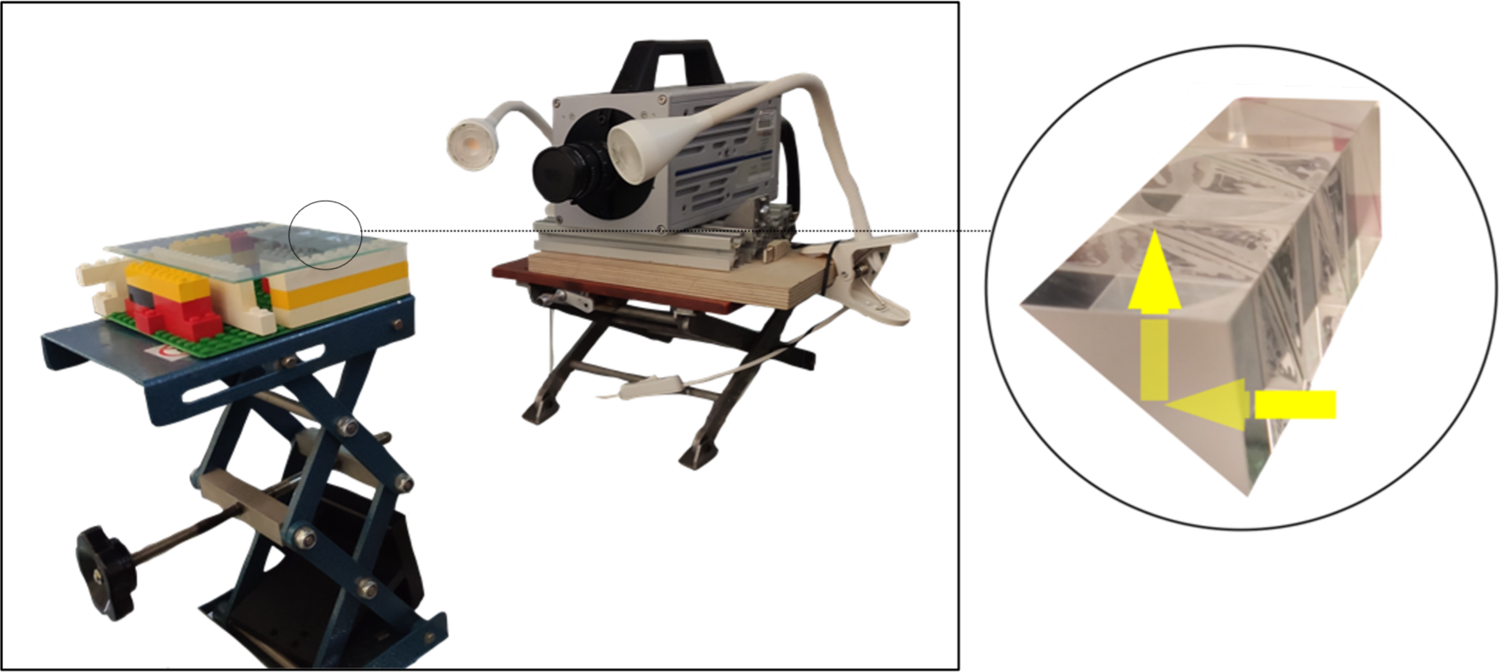
Laboratory hoist measurement area in front of the camera (left) and the row of three reflection prisms for a ventral view of the walking insects (right). Photo: Vincenz Regeler, August 10, 2022.

The frame rate was then set to 250 frames per second (fps) and the shutter speed to 1/500 seconds in the PFV 4 camera program. The frame rate was increased to 500 fps and the shutter speed reduced to 1/1000 seconds if the motion blur was too high or if the lift-off of the tarsi were not clearly recognizable. While these two settings were adjusted in the program, the light from the clamp lamps could be directed to the “region of interest” and the camera aperture iteratively adjusted so that the live image was neither too dark nor too overexposed. In addition, the trigger mode was changed to “end” so that the video recording starts to overwrite itself after a frame-rate- and resolution-dependent time window, thus minimizing “black windows” where actions of interest could have been missed. A recording was terminated with the “ready” button, whereby the buffered video could be temporarily stored and viewed under the “memory” tab. Any period of time could then be selected, named, and saved in the working directory.

**Fig. S3.**
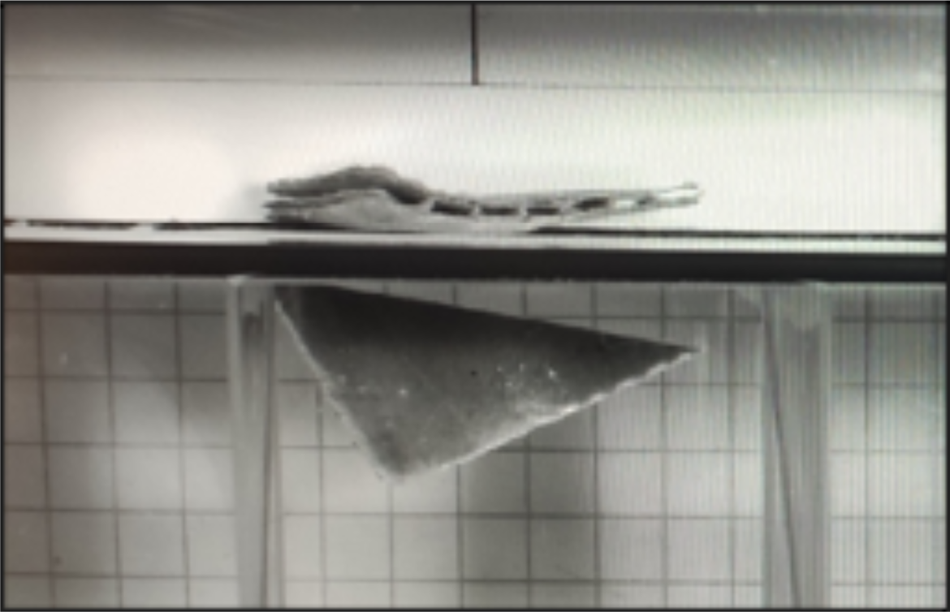
Focusing the camera with a piece of corrugated cardboard. Photo: Vincenz Regeler, August 10, 2022.

In order to ensure that the Hymenoptera could not leave the measuring area and were sufficiently constrained spatially such that they consistently walked across the “region of interest”, we constructed measuring chambers of different sizes (Fig. S4; Table S2). Their frames could be made from clamping blocks. As a transparent window for the camera, a small pane of glass was mounted on the long open side of the measuring chamber using adhesive putty, which had previously been cut to size with a glass cutter. Here it was important to ensure that the lower edge of the glass pane is flush with the entire measuring chamber so that there is no gap through which insects could escape when the measuring chamber is placed on the glass plate of the measuring area. The top of the measuring chamber could also be sealed with clamping blocks. Similar to Wöhrl *et al.,* (2017) and Reinhardt & Blickhan (2014), yet differing in scale, mathematical paper with squares of 5 mm^2^ was cut to size and glued to the inner surfaces of the measuring chambers to use as a scale for later calibration.

**Fig. S4.**
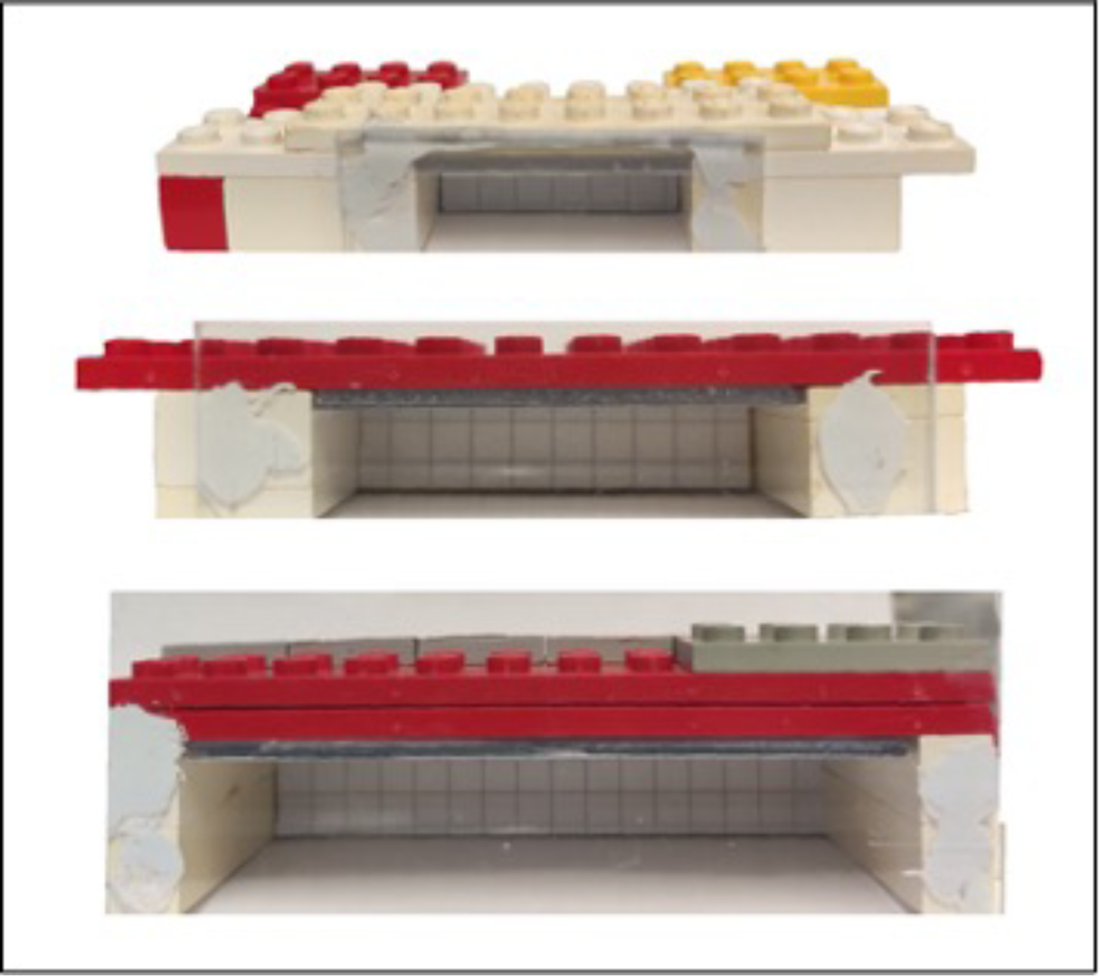
Measuring chambers in three different sizes. Photos: Vincenz Regeler, 05.09.2022.

**Table S2.** Overview of the materials of the locomotion chamber.

| Material or device designation | Manufacturer |
| --- | --- |
| Laboratory lift, DIN 12897 (stage area 160 mm x 130 mm) | Juchheim Laborgeräte GMLH; Bernkastel-Kues, Germany |
| Build Plate (127.5mm x 127.5mm) | LEGO A/S; Billund, Denmark |
| Plastic clamping blocks in various sizes:<br>- 31.8mm x 15.8mm x 9.57mm<br>- 31.8mm x 15.8mm x 3.2mm<br>- 31.8mm x 7.8mm x 9.57mm<br>- 15.8mm x 15.8mm x 9.57mm | LEGO A/S |
| Glass plate (150 mm x 105 mm) |  |
| Reflection Prism (30mm x 30mm x 30mm) |  |
| Measuring chambers in three different sizes:<br>- small (inside 30 mm x 20 mm x 15 mm)<br>- medium (inside 50 mm x 25 mm x 15 mm)<br>- large (inside 80 mm x 40 mm x 20 mm) |  |

Depending on the size of the insect to be measured, one of the three measuring chambers was selected and placed on the glass plate over the reflection prisms. The measuring chamber was then carefully lifted and the centrifuge tube containing the insect to be examined was held in so that the insect could enter the measuring chamber. It was often necessary to wait a little longer before starting the video recording the video as the insects usually groomed after handling, including wiped away drops of water from the temporary containment tubes. Once the specimen was ready, the measuring chamber could be carefully moved to ensured that the camera was positioned exactly parallel to the prisms; the video recording could then begin. As soon as the insect made an adequate locomotory lap, the recording was immediately saved with the “ready” button and viewed in order to check whether certain criteria were met. The lap itself should continue without interruption and it should be as straight as possible and centered over the prism array. More specifically, this means that the course of the lap was at a 90° angle relative to the camera, which allowed all tarsi to be seen in the lateral view (Fig. S5).

**Fig. S5.**
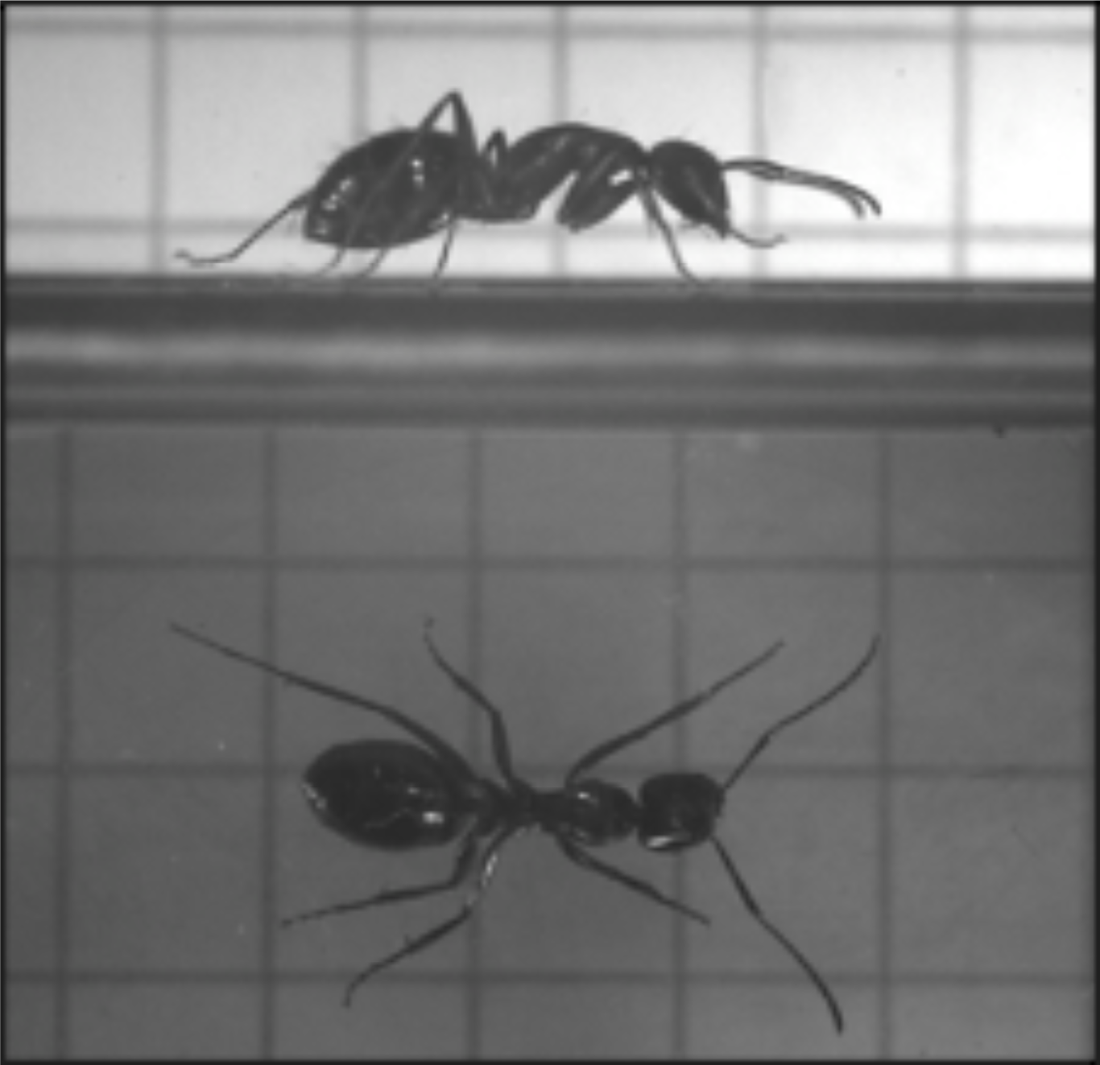
Lateral and ventral view of *Camponotus herculeanus*. Photo: Vincenz Regeler, 05.09.2022.

In addition, it was important that the legs did not touch the chamber wall in order to measure as unrestricted a run as possible on a straight surface. During the intermediate inspection, the light and contrast conditions were also checked and, if necessary, corrected so that the outlines of all tarsi could be clearly recognized. If a measurement met all quality criteria, the time period could be adjusted and saved as an MP4 or AVI file in the directory. The measurements were named according to the same pattern and numbered consecutively. For example, the file name “i09_m04” represents the fourth measurement (m04) of the ninth individual (i09).

After several measurements had been taken from an individual, the specimen was then immediately weighed using an analytical balance (Kern ABS 80-4N) (Wöhrl *et al.,* 2017). Each mass reading was recorded in milligrams (mg). The weighing always took place after the measurements, otherwise the mass before the measurement could be falsified due to the high humidity in the temporary containment tube. Deviations of up to 0.3 mg were found. Due to the limited measurement accuracy in the low milligram range (readability ± 0.1 mg, repeatability: ±0.2 mg), the analytical balance was checked at the beginning of each session using calibration weights (Fig. S6). The individuals were weighed in a sample container made of plastic and previously tared with the balance. Only light weight containers were used as the scales allowed a maximum total weight of only 83 g.

**Fig. S6.**
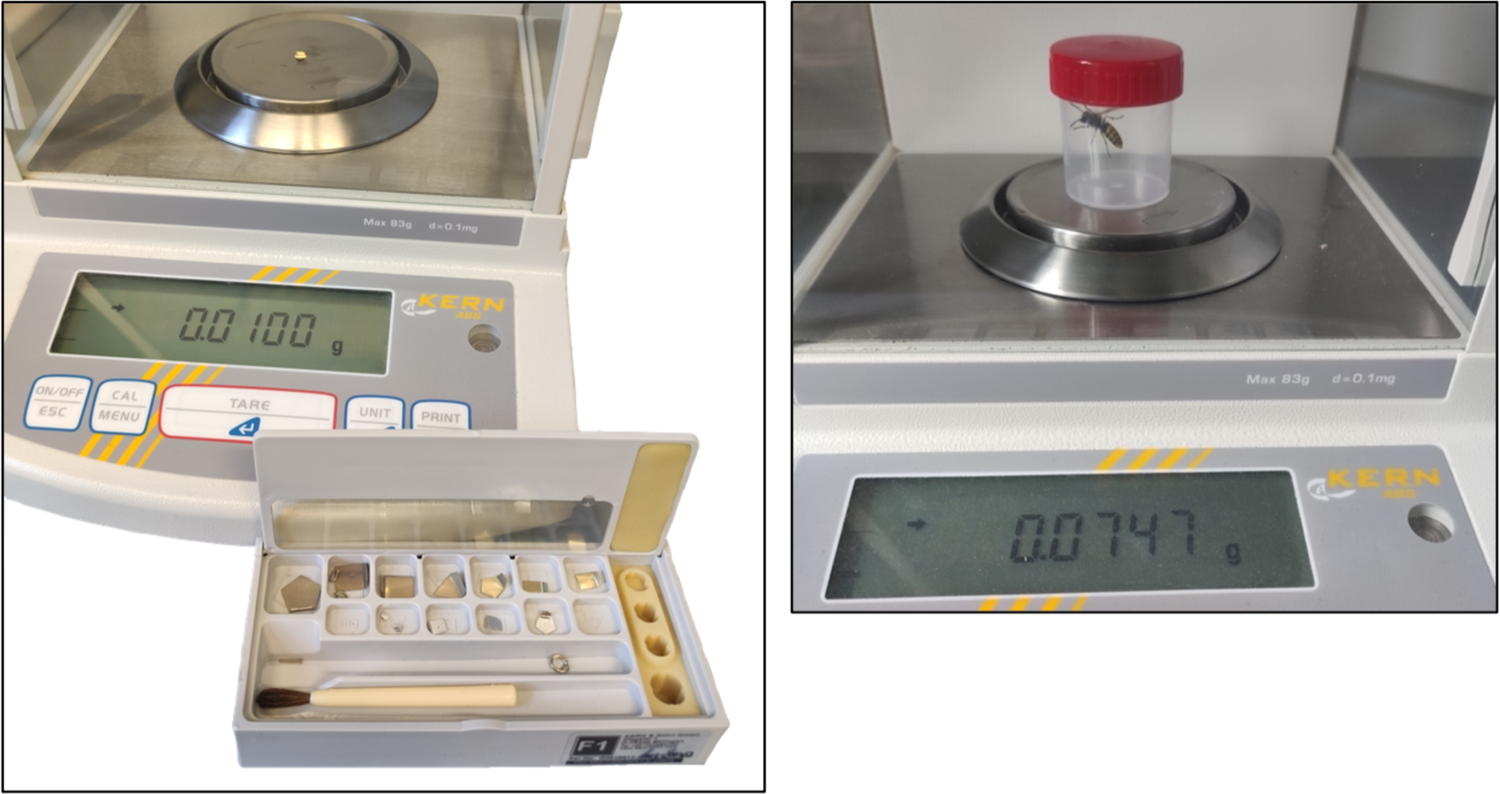
Kern ABS 80-4N analytical balance with test weights (left) and sample container (right). Photos: Vincenz Regeler, 08/04/2022.

In the case of an unequivocal identification down to the genus, the individuals were released after weighing. Otherwise, a vessel was provided with the individual number and the insect was preserved in ethanol (70%) for further determination.

#### S1.2. Tracking and data collection

A measurement number was provided for each individual that met the previously defined quality criteria. Further processing of these selected measurements was initially carried out in the program Tracker, which is a free video analysis program and modeling tool based on the opensource Physics Java Library (OSP) (Brown *et al.,* 2022). Among other things, the program allows the user to zoom in on video files, for which reason we used it between the measurements to check the camera recording for contrast, brightness, and the effect of motion blur.

The video files (MP4 / AVI) were opened in the Tracker program; the frame rate of each recording had to be individually specified. Then, with the help of the box scale in the video, a calibration standard could be set, and its dimensions determined (Fig. S7). This allowed the image coordinates (pixels) to be converted into a unit of length. In the next step, eleven different point masses (“trackers”) were defined (Table 1 in main text), and their positions were added manually frame by frame with a mouse click (Fig. S8). Selecting a position automatically loads the next frame. The different point masses were tracked on up to 320 consecutive images, so that each measurement enabled the investigation of three stepping cycles (Wahl *et al.,* 2015). Up to 3,500 positions had to be added manually in one measurement.

**Fig. S7.**
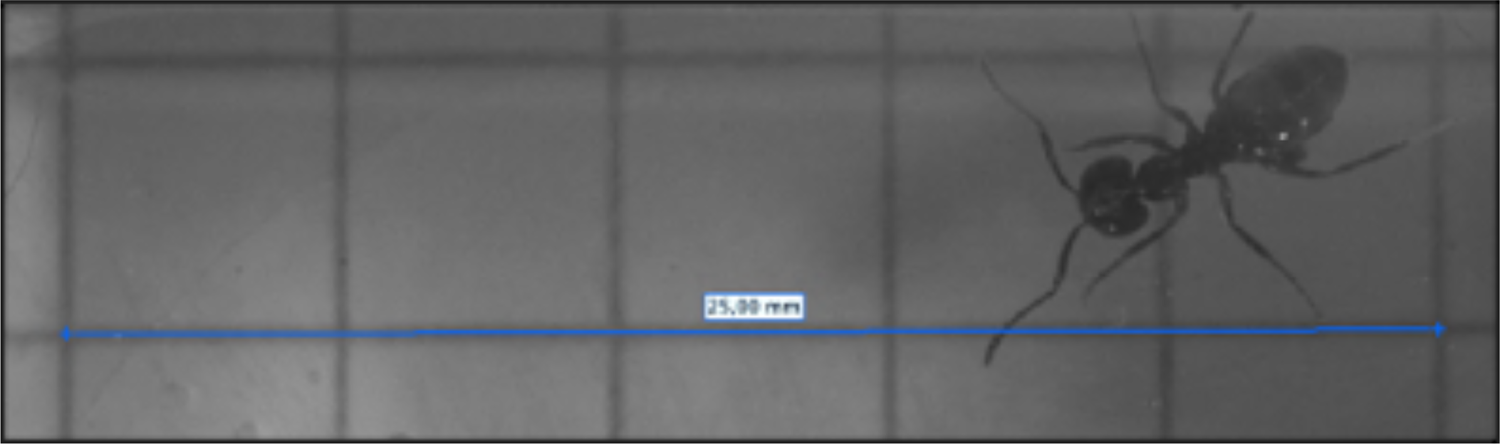
Defining the calibration scale on the mathematical paper attached to the roof of the chamber.

All point masses were set at the same time from the beginning of a swing phase of the left front leg. The condition for this was that two other clearly recognizable swing phases of the same leg were recorded in the video.

Unlike tarsi, coxae were only tracked on the left side of the leg—unless the right side of the leg was more discernible or less affected when running. Under these circumstances, a right swing phase was started and only the right coxae were tracked. After completion, the processing log of a video file could be saved as an XML-based tracker file (.trk), which contains the information about the set step positions as image coordinates. However, before the tracker files went into further processing, the videos were double-checked in the program in order to determine the temporal step pattern.

**Fig. S8.**
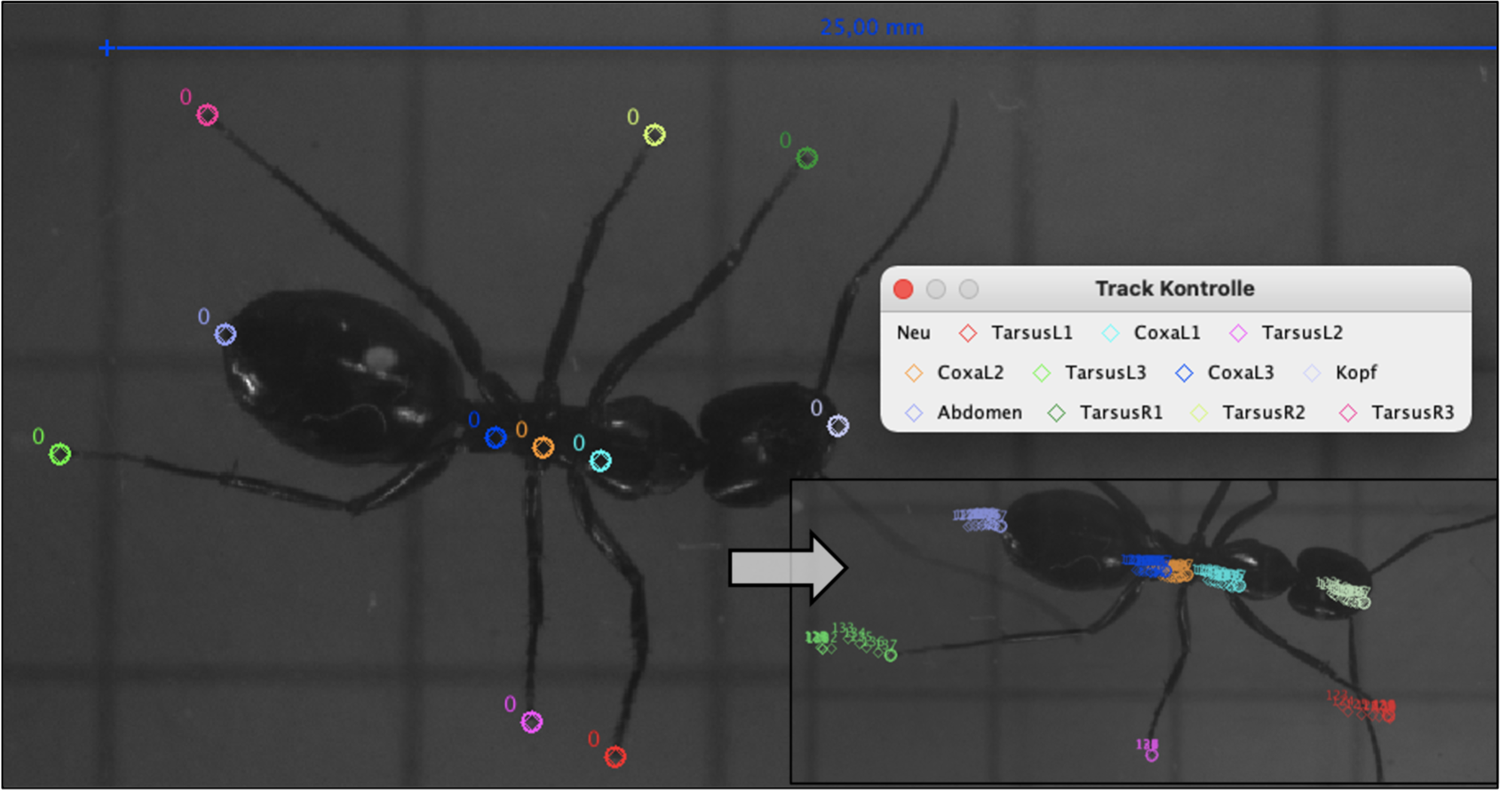
Ventral view of *Camponotus herculeanus* with set positions.

#### S1.3. Analysis of temporal stepping pattern

For each video, a new start image could be set to the point in time when the point masses were first added. Starting from this start image, the videos were examined in a similar way to the work by Humeau *et al.,* (2019). The movement of all tarsi was observed frame by frame with the naked eye. For each tarsus, the image numbers of lifting (“lift-off”) and planting (“touchdown”) could be noted over three swing phases and recorded in an Excel spreadsheet (Fig. S9). According to this, 36 image numbers were recorded as points in time for each measurement, and two complete contact phases could also be derived from them. Since some Hymenoptera dragged their hind legs over the ground, attention was paid to the onset of an active movement in the metatarsi, and as soon as the tarsus left its original contact point, the swing phase was considered to start (lift-off time of Reinhardt & Blickhan 2014, Wahl *et al.,* 2015).

**Fig. S9.**
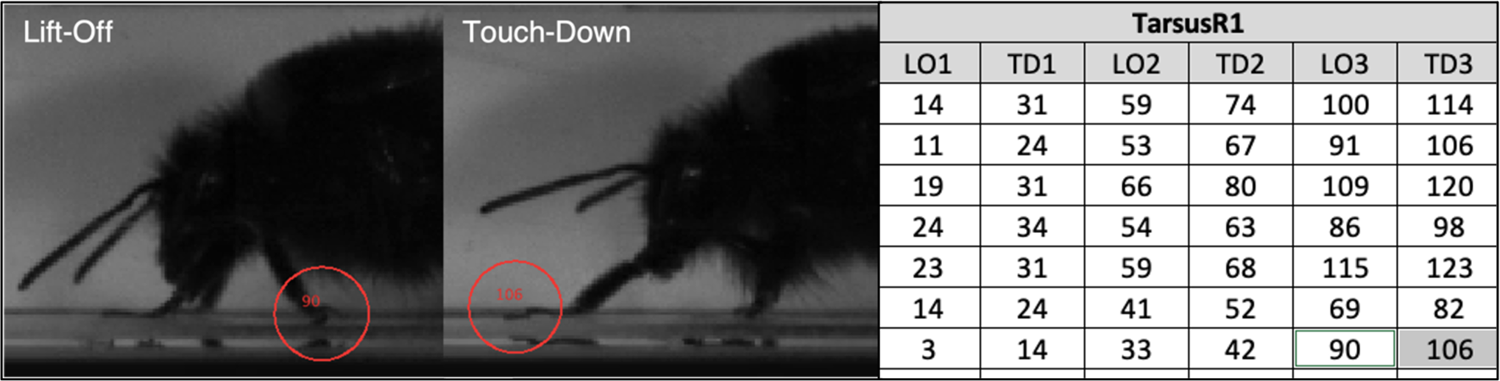
Lifting and landing times of the right protarsus (TarsusR1) (left). Section of the Excel table with image numbers for the liftoffs (LO) and touchdowns (TD) (right).

After completing the table, the readings were converted to seconds depending on their frame rate. To do this, the values had to be divided by 250 or 500, depending on the frame rate. For example, 10 frames captured at a frame rate of 250 fps gives a time of 0.04 seconds (10 ÷ 250 = 0.04).

For each individual, a separate Excel workbook was created, into which a previously created template was inserted. This included table cells with calculation functions for the duration of the swing and contact phases, which were also linked to the creation of a stacked bar chart. In order to have Excel calculate the swing and contact times, only the times when the tarsi lifted and touched down had to be entered in the area with the green-painted background (Fig. S10).

**Fig. S10.**
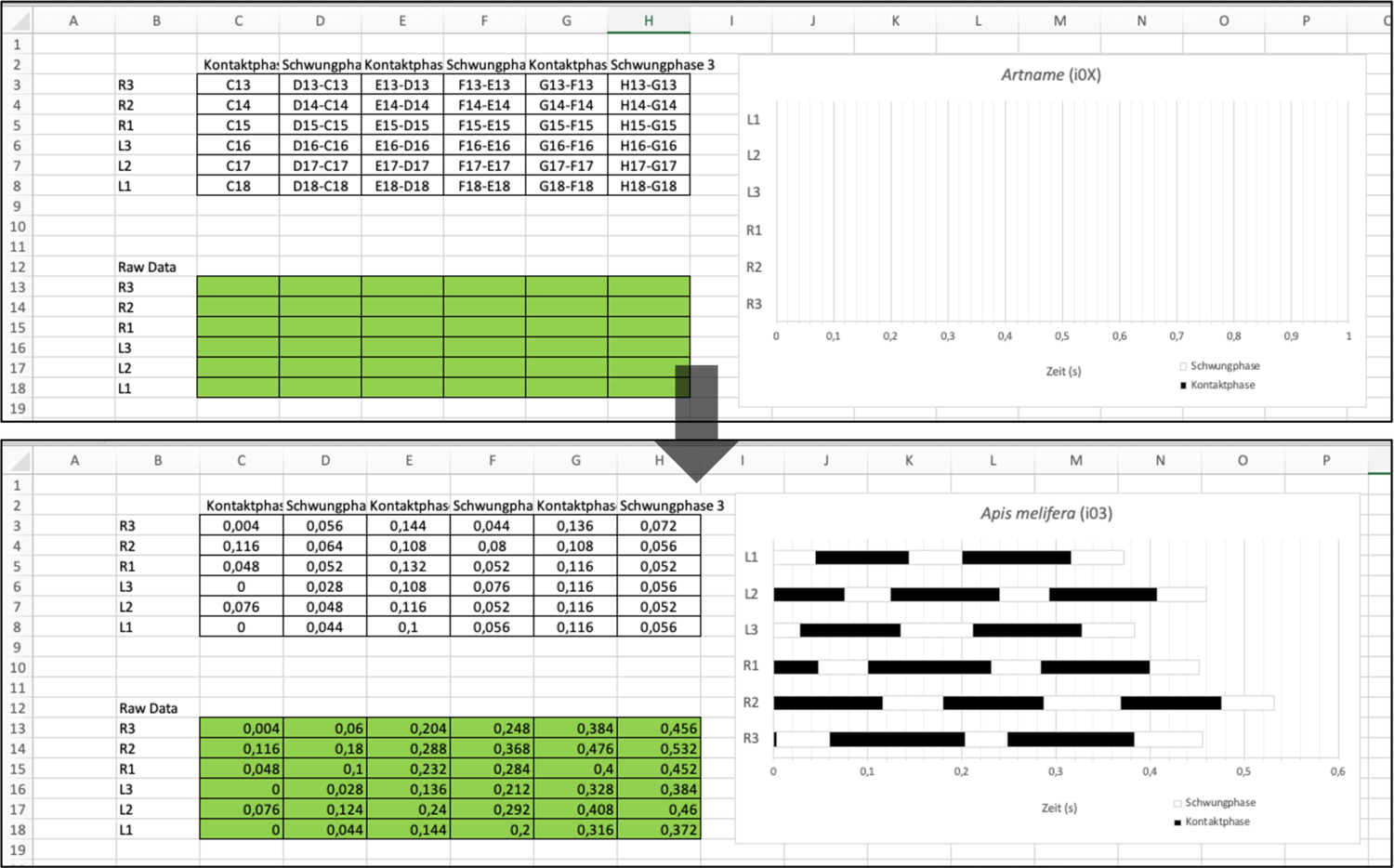
Excel template with calculation functions for creating the temporal step pattern (above) and an inserted example for *Apis mellifera* (below).

Now the swing phases were colored white and the contact phases black (Zollikofer 1994a; Grabowska *et al.,* 2012). The chart was titled, the legs numbered on the Y-axis, and the time in seconds on the X-axis. Ultimately, the temporal step patterns could be saved in PDF format. This type of representation is a traditional, established tool in temporal step pattern analysis. For example, as early as 19^th^ century, the pace and gallop pattern of horses could be represented in a similar way (Marey 1874).

For reasons of comparability, another diagram template was created, which shows the step pattern in a simplified way and scaled with a total time span of one second (Fig. S11). In addition, the mean values of the measurements were used to create the temporal step pattern at the systematic family and superfamily level. Assuming that the left and right strides are more or less the same, the stride patterns may have been mirrored so that all patterns begin with a left front leg (L1) swing phase as a reference point.

**Fig. S11.**
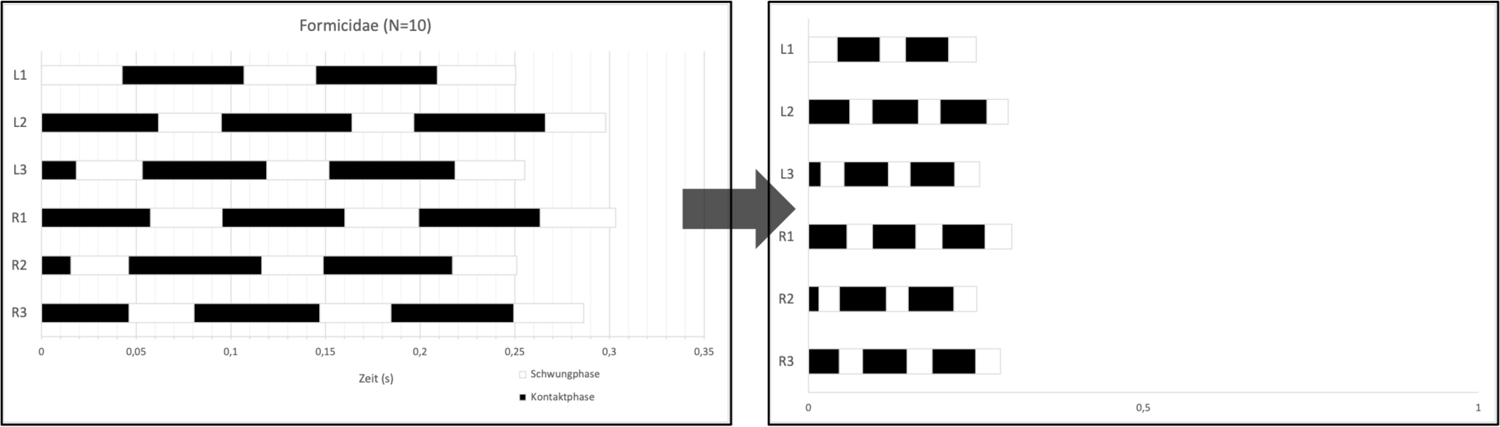
The conversion of a temporal stepping pattern into a simplified, scaling pattern.

The duty factor was calculated from the relationship between swing time and contact time (Weihmann *et al.,* 2017; Wahl *et al.,* 2015). The formula for calculating the duty factor was as follows:

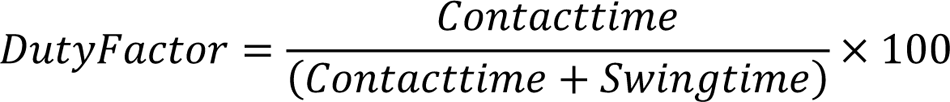

#### S1.4. Analysis of spatial stepping pattern

In order to determine the spatial step patterns of the sampled individuals, the area to which the point masses were added was viewed again in the Tracker program. From this, three individual images were noted for each measurement, at which times three legs were in a contact phase and the other three legs were in a swing phase (Zhao *et al.,* 2018). This had to be the middle leg on one side of the body and the front and rear legs on the other side of the body (tripod gait), so that if the positions of the point masses were connected, a triangle would result (Fig. S12). The times were determined in three consecutive steps, so that the tripodal leg groups alternated (Zollikofer 1994a; Seidl & Wehner 2008; Wahl et al., 2015; Pfeffer et al., 2019).

**Fig. S12.**
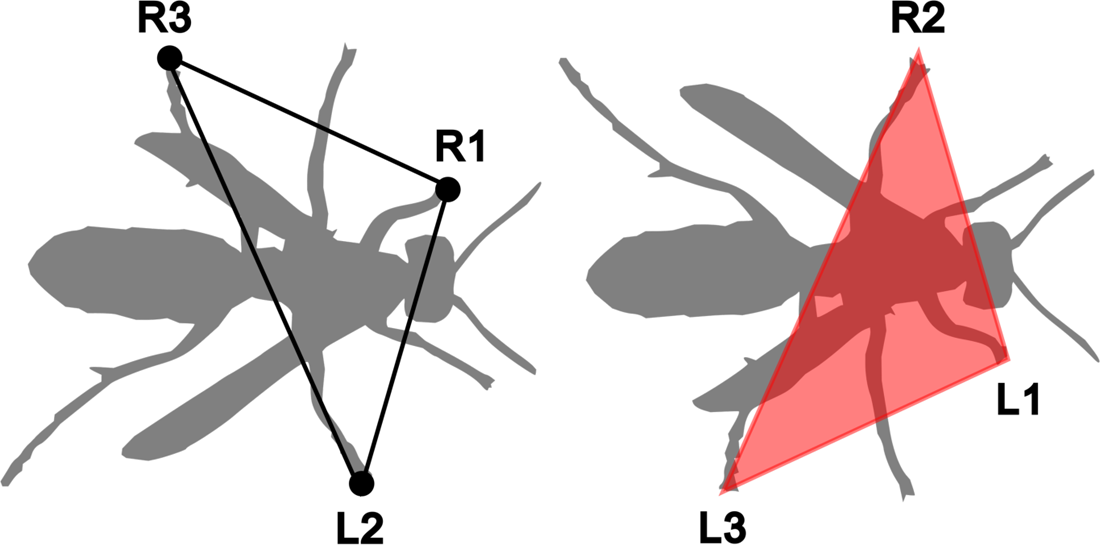
Two consecutive tripod stances, exemplified by *Polistes* sp.

The point in time at which the middle leg in the swing phase overtook the standing middle leg was selected as a point of reference for the determination of the three individual images. At the same time, the image numbers of the tripod individual images were recorded in an Excel spreadsheet.

A project was then created in RStudio and an R markdown file (.rmd) was created therein. This makes it possible to combine text and code from the R programming language. The R markdown file could then be converted to an HTML file, in which the program code is executed, and the associated text is also displayed. With a function created in the program code, the data could be imported from the tracker files. This data contains information about the calibration scale and about the point masses added in the frames and their pixel coordinates. Using various commands in R, calculations could be carried out and points plotted, *i.e.*, displayed graphically. The correct and consistent naming of the point masses was essential to reduce errors in the program code as well as to speed up and automate the process.

The coordinates of the relevant tarsi (L1, R2, L3 or R1, L2, R3) from the respective three individual images were connected to form triangles. The point masses for the head and abdomen could be plotted as a connected line to serve as a reference for the body-fixed coordinate system. The lengths of the individual lines and the distances between the point masses could also be calculated. This made it possible to determine body length, stride length, and step width. The middle between the point masses of head and abdomen represents the central point. The distance covered by this point was calculated for the stride length, in the time span from the touch-down of the middle tarsus to the next touchdown of the same tarsus (Zollikofer 1994a; Wahl *et al.,* 2015; Pfeffer *et al.,* 2019). The step width indicates the lateral distance from the center of the body to the touchdown point of the tarsus in millimeters and is made up of the mean values of a pair of legs (L1, R1; L2, R2; L3, R3).

#### S1.5. Analysis of range of motion

The most proximal segment (coxa) and the most distal segment (tarsus) were chosen to determine the range of motion of the legs. For simplification, it was assumed that the insect leg moves more or less in one plane (“leg plane”), since the coxa-trochanter, femur-tibia, and tibia-tarsus joints are joints aligned in parallel (Cruse & Bartling 1995; Seidl & Wehner 2008; although see Boudinot 2015 for the procoxotrochanteral articulation of Formicidae and Arroyave-Tobon *et al*. 2022 for the trochanterofemoral joint). From the ventral view, the angle from the longitudinal body axis (BLA) to the anterior (AEP) and posterior (PEP) extreme position of the tarsus and the angle between them could be calculated from the coxa.

The “minimum” angle is between the BLA and AEP, and the “maximum” angle is between the BLA and PEP (Fig. S13). The variation in leg movement means that the AEPs and PEPs in the body coordinate system always change slightly (Wosnitza *et al.,* 2013), for which reason the range of motion could only be determined over several steps. As with the spatial stepping pattern, the point masses of head and abdomen were used to establish the body-fixed coordinate system. This now made it possible to plot the coxae with the associated tarsus across the individual images with connecting lines (CoxaL1, TarsusL1; CoxaL2, TarsusL2; CoxaL3, TarsusL3 or analogously for the right side of the leg). Ultimately, the resulting “*Schneeengel*” plots (trajectories) of the individuals were all scaled to the same size, mirrored to the left side of the body if necessary and, for visual understanding, combined with a manufactured contour of the individual as described by Wöhrl *et al*. (2017).

**Fig. S13.**
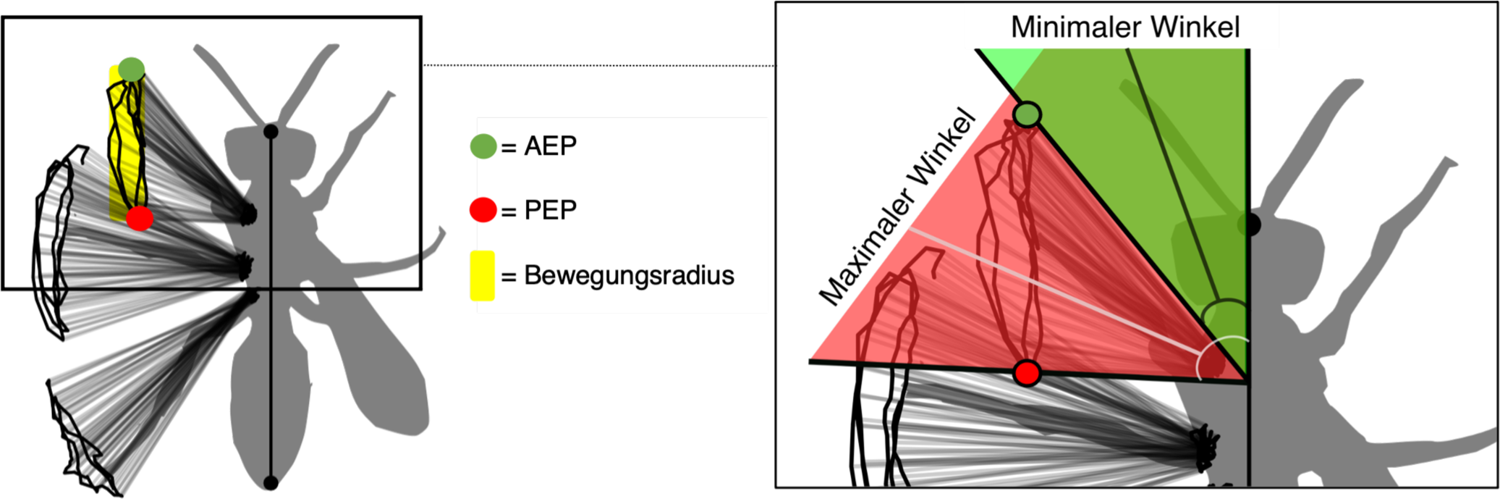
Trajectories of the tarsi, shown as a *Schneeengel* pattern with contour, using the example of *Cerceris* sp. including the AEP and PEP of the protarsus as well as the representation of the minimum and maximum angle.

To create the contours, a screen shot of the individual was taken from the respective measurement. These were edited in the MacOS program “Preview”. With the lasso tool, the outlines of the Hymenoptera could be traced and cut out by dragging the mouse. This step was sometimes made more difficult by the fact that the videos were too low in brightness and/or image quality (Fig. S14). Outlines with and without wings were created for flying Hymenoptera.

**Fig. S14.**
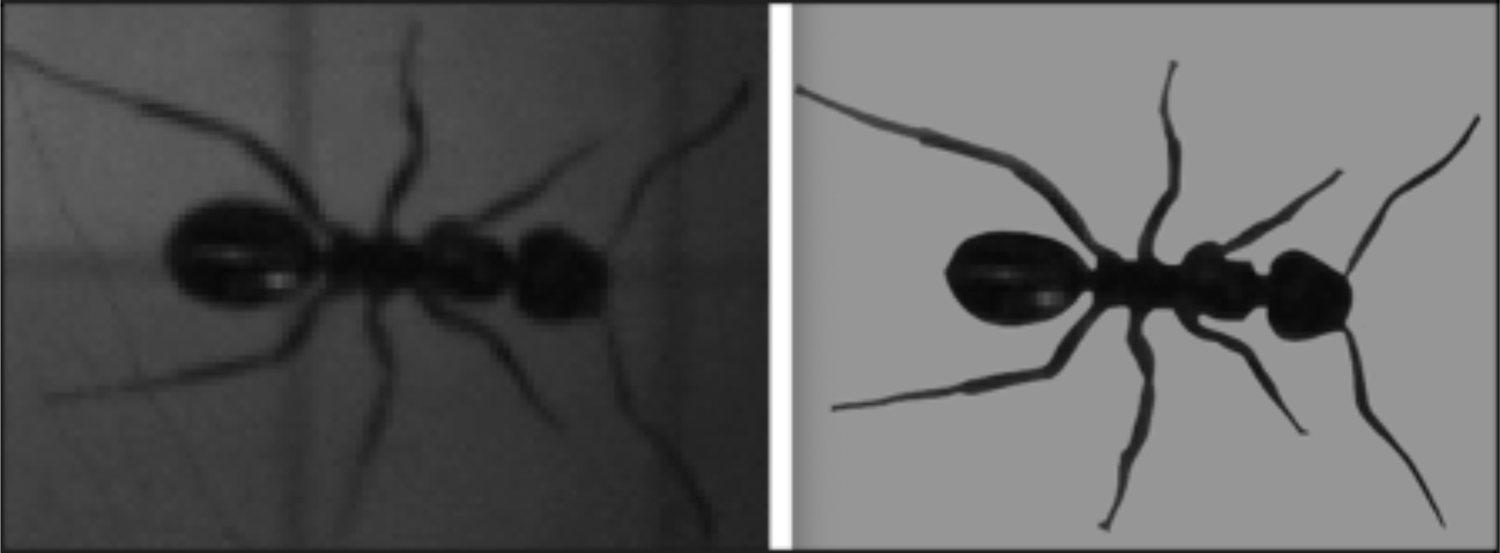
Screen capture (left) and cropped outline (right) of *Formica* sp.

The clipped outlines were then editable in PNG format in the free Inkscape drawing program. Here, the respective contour was created with the “Trace bitmap” option and a brightness thresh-old of 0.95 and then colored gray (40%). The *Schneeengel* illustrations were inserted into Inkscape as a PDF file. After that, the contours could be moved one level behind the *Schneeengel* plots and adjusted to their height scale and aligned accordingly. For this, the same positions were chosen as when setting the point masses (head and abdomen) and not the end of the wings or the ovipositor.

## S2. Additional Results & Discussion

In this supplementary section, we provide greater detail for select results (S2.1) and we report comparisons based on ecology and certain behavioral features (S2.2), for which there is inadequate space in the main text. Supplementary results figures relating to the main text are presented at the end of S2.1 and additional data tables relating to the main text are provided at the end of S2.2, which also includes figures specific to this section.

### S2.1. Further detail for results in main text

#### S2.1.1. Results aggregated at superfamily level

As the phylogeny of Hymenoptera is largely resolved at the family level (Peters *et al*. 2017; Branstetter *et al*. 2017), the superfamily rank provides a monophyletic aggregation unit for our sampled taxa. At this level of comparison, an apparent phylogenetic trend in temporal gait pattern from the most recent common ancestor of the order to the Formicidae is visible (Fig. S15), with sawflies (Tenthredinoidea) having low step frequencies and long contact times, the non-aculeate Apocrita (Ichneumonoidea, Cynipoidea, Evanioidea) having higher frequencies and shorter contact times, and finally through the Aculeata, the Formicidae having the greatest frequency and shortest contact times. Additional observations aggregated at this rank are provided in Fig. 4. Most critically for the deeper evolution of the Hymenoptera, we observe that the sampled sawflies have exceptionally wide foreleg tracks (Fig. 4, Char. 8) and never displayed a tripodal gait (Fig. 4, Char. 1). As discussed in the main text, the character polarity of these conditions remains uncertain without expanded sampling of sawfly superfamilies. Such sampling will also allow for quantitative phylogenetic reconstruction of the results.

**Fig. S15.**
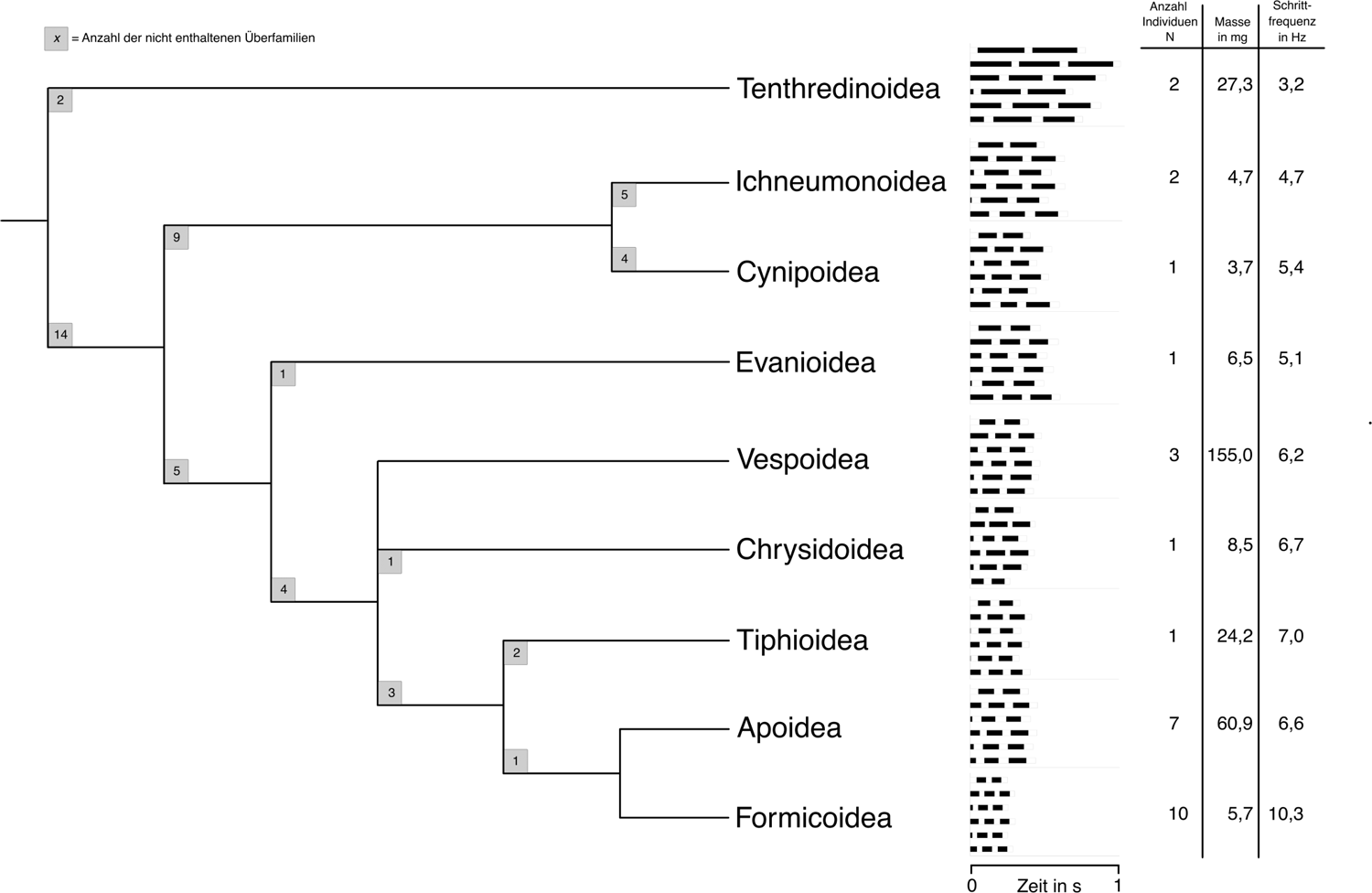
Cladogram of the examined Hymenoptera superfamilies including average temporal step pattern, number of individuals N, average mass in mg, and average step frequency in Hz.

#### S2.1.2. Results aggregated at family to genus level

Within the monophyletic superfamilies, our sampling was variable at the genus to family rank, thus we provide results aggregated at these levels more-or-less interchangeably (Figs. S16, S17). Because kinematic studies of hymenopteran locomotion are restricted to *Apis mellifera* and Formicidae, we discuss five notable taxa here: *Gasteruption* (Gasteruptiidae: Evanioidea), *Tiphia* (Tiphiidae: Tiphioidea), *Vespa* (Vespidae: Vespoidea), and the two sampled ichneumonoid genera.

**Fig. S16a.**
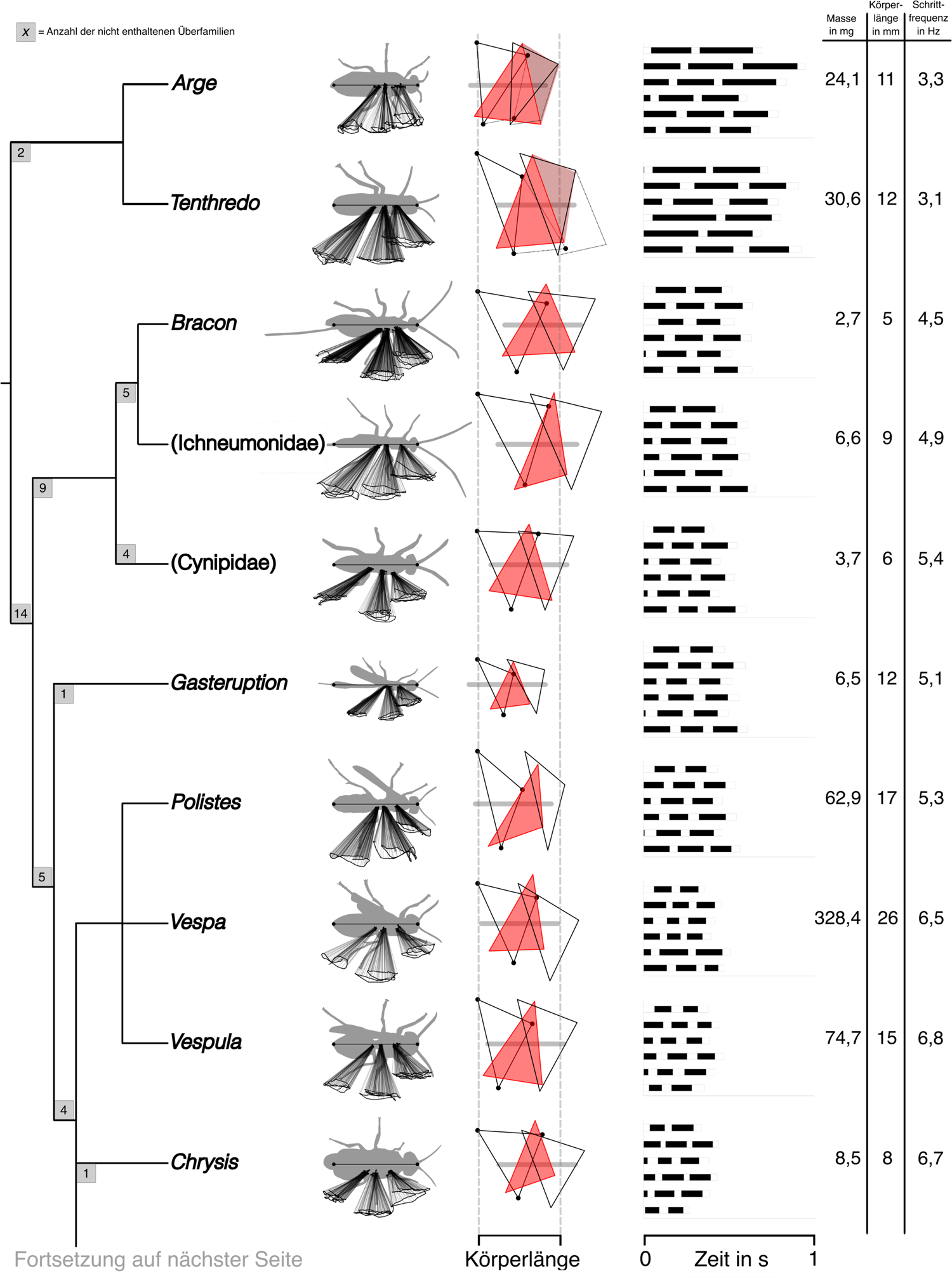
Phylogeny of the examined genera with contours and *Schneeengel* patterns as well as spatial and temporal stepping patterns and information columns for mass in mg, body length in mm and stepping frequency in Hz. Contours in ventral view, running direction from left to right. (1/2). Topology from Peters *et al*. (2017) and Branstetter *et al*. (2017).

**Fig. S16b.**
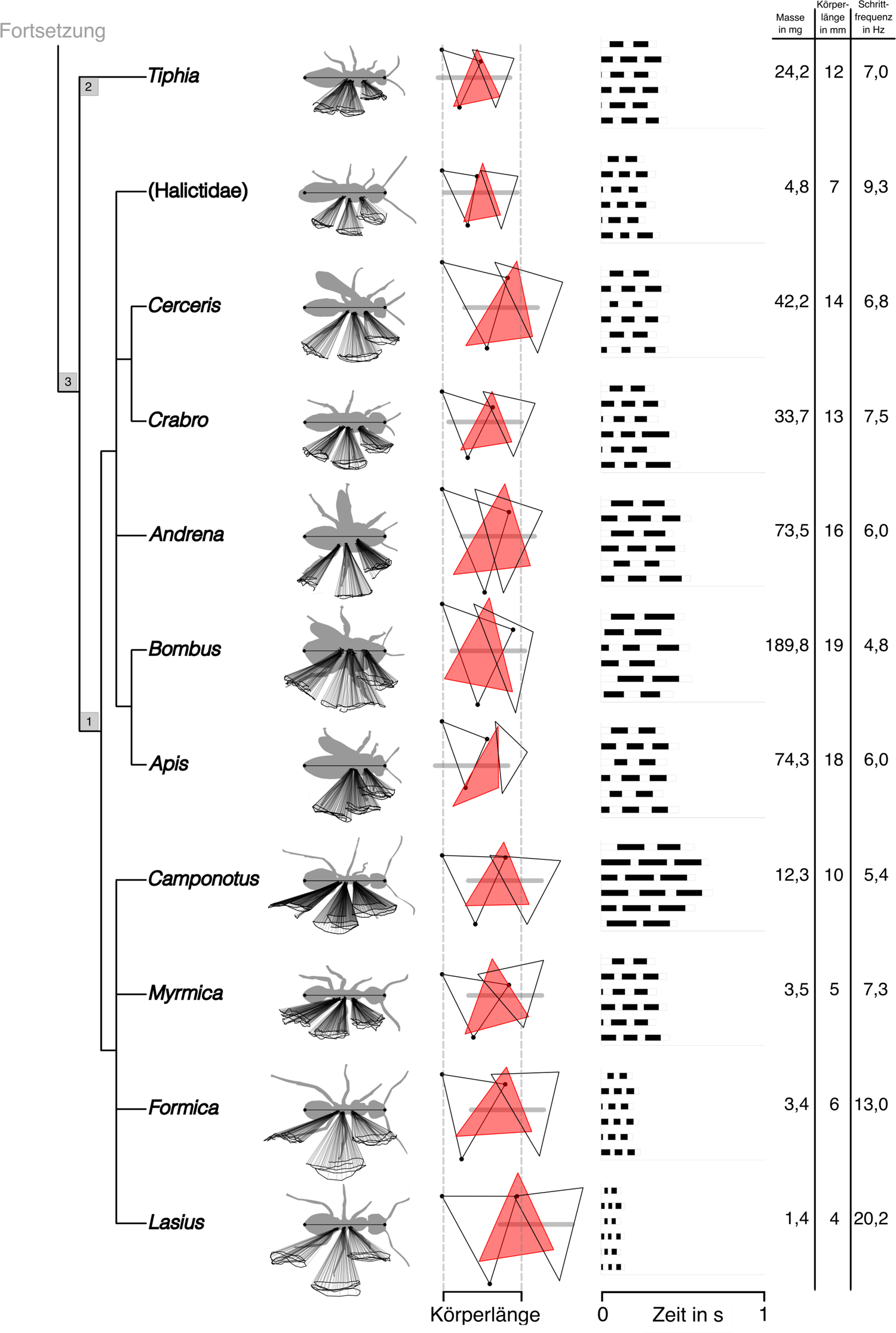
Phylogeny of the examined genera with contours and *Schneeengel* patterns as well as spatial and temporal stepping patterns and information columns for mass in mg, body length in mm and stepping frequency in Hz. Contours in ventral view, running direction from left to right. (2/2).

**Fig. S17.**
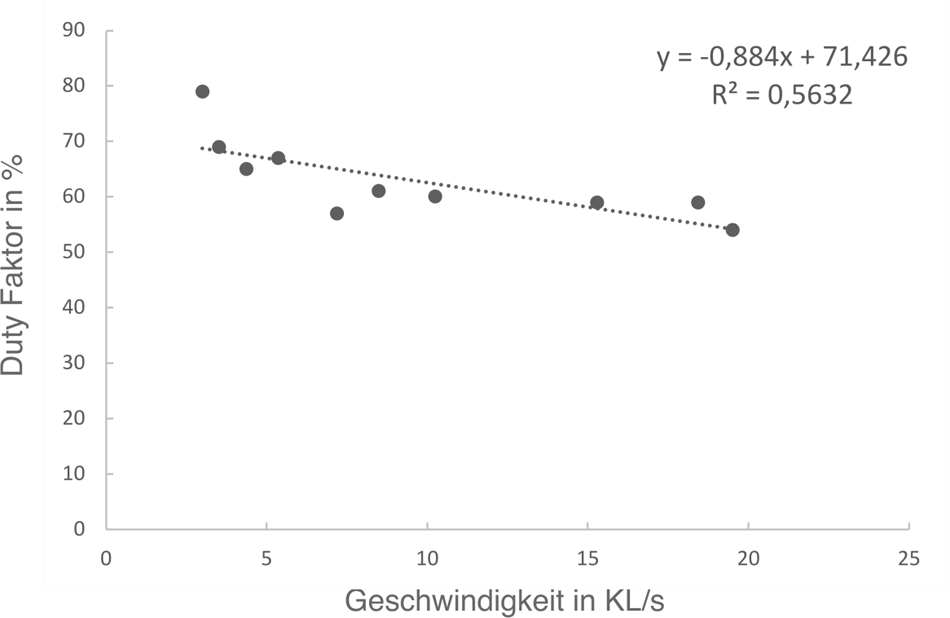
Ratio of the duty factor in % to the relative speed in body lengths per second (BL/s) from the mean values of all examined ants (N = 10). Linear regression is shown as a dotted line.

We observed that the sampled *Gasteruption* had limited walking capacity. The individual moved at a low speed of 1.9 BL/s and had a low TCS value of 0.1 (Table S6), and it frequently toppled over having lost safe contact with the flooring surface of the measurement chamber. The latter can be reasonably attributed to the disproportionately long and posteriorly thickened metasoma of the genus, which was dragged throughout the sampling period, although the unstudied attachment structures and the fat-filled metatibiae (Mikó *et al.,* 2019) may also play a role. We find the toppling and dragging to be of special interest given the extremely dorsal and anterior positioning of metasomal-propodeal articulation, nearly abutting the metanotum; the consequences of this derived articulation are poorly if at all understood. It is possible that Gasteruptiidae and its sister family, Aulacidae, may behave like odonates when walking, either gripping onto very rough surfaces or onto rounded stems and narrow leaves. *Gasteruption* also achieved the widest range of motion angle in the forelimbs (82°) and the lowest angle in the middle legs (51°) (Fig. S16a; Table S7). The short legs, already known to be a limiting factor (*e.g.*, Reinhardt & Blickhan 2014; Nirody 2021; Wittlinger et al., 2007; Wahl et al., 2015), can reasonably explain the short stride length and narrow step width.

**Fig. S18.**
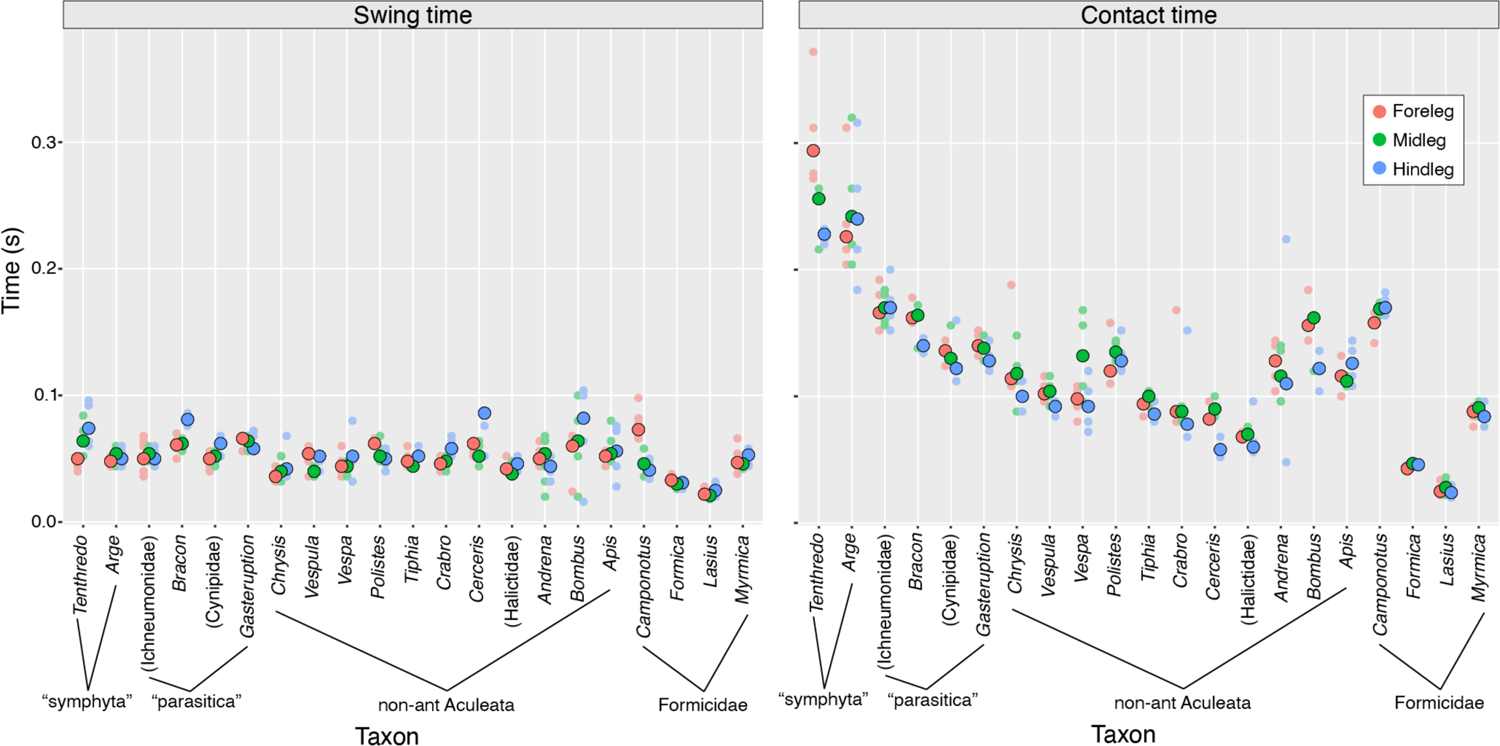
Swing and contact times for the sampled Hymenoptera (each leg n = 6), aggregated by genus. Both panels to same scale. Large, opaque dots indicate median values per leg; smaller, transparent dots plot all observations. Where generic identifications unavailable, family indicated in parentheses. Brackets indicate groupings of the sampled taxa.

**Table S3.** Overview of other materials.

| Material or device designation | Manufacturer |
| --- | --- |
| Analytical balance Kern ABS 80-4N | Kern & Sohn GMLH; Balingen, Germany |
| Fractional case with F1 fine/test weights | Kern & Sohn GMLH |
| Plastic sample container with lid |  |
| 15mL centrifuge tube |  |
| 25 ml snap-top jar (Ø 30 mm) |  |

**Table S4.**
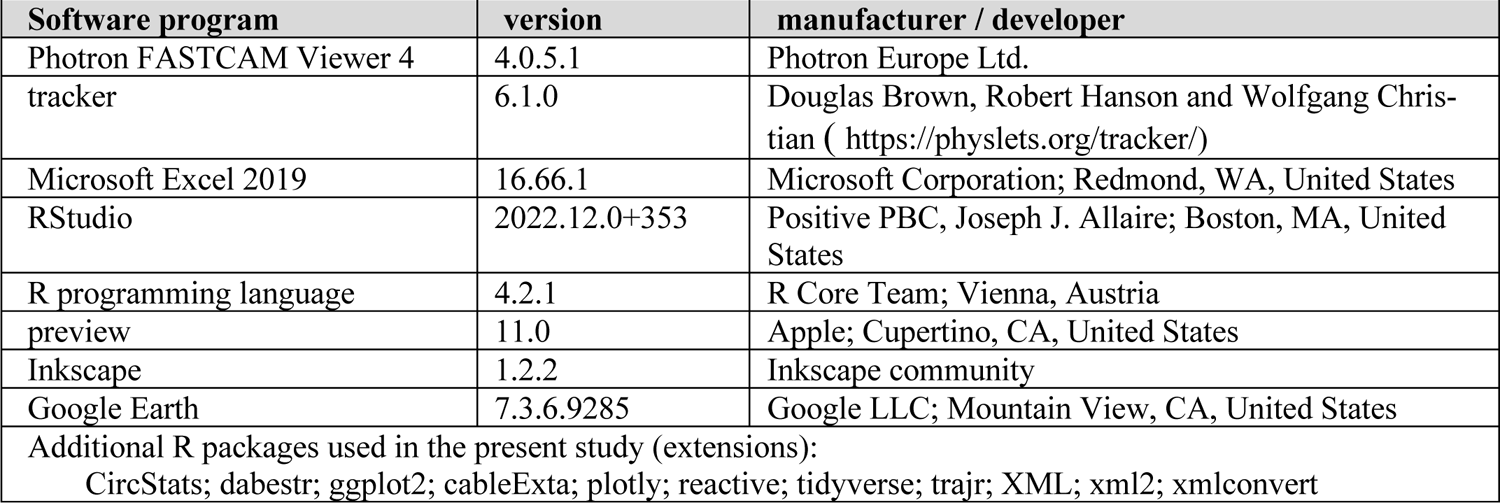
Overview of the software programs used.

**Table S5.** Overview of the individuals collected, including their geographic coordinates, microhabitat, and collector information. Collector abbreviations: BEB = Dr. Brendon E Boudinot; VR = Vincenz Regeler. The grayed-out rows were not included in the locomotory analyses as the individual served as a pre-measurement or belongs to another insect order.

| ID | Order | Superfamily | Family | Genus species | Collecting information |
| --- | --- | --- | --- | --- | --- |
| 1 | Hymenoptera | Apoidea | Apidae | *Apis mellifera*<br>Linnaeus, 1758 | 50°43'04.4"N 11°20'18.4"E,<br>common viper's bugloss. VR |
| 2 | Hymenoptera | Formicoidea | Formicidae | *Lasius* sp. | 50°43'04.4"N 11°20'18.4"E<br>gravel bed. VR |
| 3 | Hymenoptera | Apoidea | Apidae | *Apis mellifera* | 50°55'21.0"N 11°35'06.7"E<br>common loosestrife. VR |
| 4 | Hymenoptera | Vespoidea | Vespidae | *Vespa germanica*<br>(Fabricius, 1793) | 50°55'27.9"N 11°35'00.9"E<br>wild cherry, on fruits. VR |
| 5 | Hymenoptera | Apoidea | Andrenidae | *Andrena* sp. | 50°55'21.0"N 11°35'06.7"E<br>common loosestrife. VR |
| 6 | Hymenoptera | Vespoidea | Vespidae | *Vespa crabro*<br>Linnaeus, 1758 | 50°55'27.9"N 11°35'00.9"E<br>wild cherry. BEB |
| 7 | Hymenoptera | Chrysidoidea | Chrysididae | *Chrysis* sp. | 50°55'27.9"N 11°35'00.9"E<br>wild cherry. BEB |
| 8 | Hymenoptera | Apoidea | Halictidae | (Halictidae) | 50°55'27.9"N 11°35'00.9"E<br>wild cherry. BEB |
| 9 | Hymenoptera | Apoidea | Apidae | *Bombus terrestris*<br>(Linnaeus, 1758) | 50°55'24.0"N 11°35'00.6"E<br>ordinary snowberry. VR |
| 10 | Diptera | Oestroidea | Tachinidae | *Tachina fera*<br>(Linnaeus, 1761) | 50°55'24.0"N 11°35'00.6"E<br>ordinary snowberry. VR |
| 11 | Hymenoptera | Tenthredinoidea | Tenthredinidae | *Tenthredo* sp. | 50°55'11.8"N 11°35'00.9"E<br>wild carrot. BEB |
| 12 | Hymenoptera | Tenthredinoidea | Argidae | *Arge* sp. | 50°55'11.8"N 11°35'00.9"E<br>wild carrot. BEB |
| 13 | Hymenoptera | Tiphioidea | Tiphiidae | *Tiphia* sp. | 50°55'11.8"N 11°35'00.9"E<br>wild carrot. BEB |
| 14 | Hymenoptera | Apoidea | Crabronidae | *Cerceris* sp. | 50°55'11.8"N 11°35'00.9"E<br>wild carrot. BEB |
| 15 | Hymenoptera | Evanioidea | Gasteruptiidae | *Gasteruption* sp. | 50°55'11.8"N 11°35'00.9"E<br>wild carrot. BEB |
| 16 | Strepsiptera | - | - | - | Collector: Dr. Hans Pohl |
| 17 | Hymenoptera | Cynipoidea | Cynipidae | (Cynipidae) | 50°55'11.8"N 11°35'00.9"E<br>wild carrot. BEB |
| 18 | Hymenoptera | Ichneumonoidea | Ichneumonidae | (Ichneumonidae) | 50°55'11.8"N 11°35'00.9"E<br>wild carrot. BEB |
| 19 | Hymenoptera | Apoidea | Crabronidae | *Crabro* sp. | 50°55'16.0"N 11°34'57.9"E<br>Sandy bottom. BEB |
| 20 | Hymenoptera | Apoidea | Crabronidae | *Cerceris* sp. | 50°55'16.0"N 11°34'57.9"E<br>Sandy bottom. BEB |
| 21 | Hymenoptera | Formicoidea | Formicidae | *Lasius fuliginosus*<br>(Latreille, 1798) | 50°55'21.2"N 11°35'06.3"E<br>wayside. VR |
| 22 | Hymenoptera | Ichneumonoidea | Braconidae | *Bracon* sp. | 50°55'24.0"N 11°35'00.6"E<br>grapevine. VR |
| 23 | Hymenoptera | Vespoidea | Vespidae | *Polistes* sp. | 50°55'11.8"N 11°35'00.9"E<br>wild carrot. BEB |
| 24 | Hemiptera | Pyrrhocoroidea | Pyrrhocoridae | *Pyrrhocoris apterus*<br>(Linnaeus, 1758) | 50°55'27.9"N 11°35'00.9"E<br>wild cherry. BEB |
| 25 | Hemiptera | Pentatomoidea | Pentatomidae | - | 50°55'27.9"N 11°35'00.9"E<br>wild cherry. BEB |
| 26 | Hymenoptera | Formicoidea | Formicidae | *Formica* cf. *fusca* | 50°55'27.9"N 11°35'00.9"E<br>sandy bottom. VR |
| 27 | Hymenoptera | Formicoidea | Formicidae | *Lasius niger*<br>(Linnaeus, 1758) | 50°55'27.9"N 11°35'00.9"E<br>colony in the sand. BEB |
| 28 | Hymenoptera | Formicoidea | Formicidae | *Lasius niger* | 50°55'27.9"N 11°35'00.9"E<br>sandy bottom. VR |
| 29 | Hymenoptera | Formicoidea | Formicidae | *Myrmica* sp. | 50°55'27.9"N 11°35'00.9"E<br>sandy bottom. VR |
| 30 | Hymenoptera | Formicoidea | Formicidae | *Myrmica* sp. | 50°55'27.9"N 11°35'00.9"E<br>sandy bottom. VR |
| 31 | Neuroptera | - | - | - | Collector: Di Li |
| 32 | Neuroptera | - | - | - | Collector: Di Li |
| 33 | Neuroptera | - | - | - | Collector: Di Li |
| 34 | Neuroptera | - | - | - | Collector: Di Li |
| 35 | Hymenoptera | Formicoidea | Formicidae | *Lasius* sp. | 50°55'10.6"N 11°35'00.3"E<br>wayside. VR |
| 36 | Hymenoptera | Formicoidea | Formicidae | *Lasius* sp. | 50°55'10.6"N 11°35'00.3"E<br>wayside. VR |
| 37 | Hymenoptera | Formicoidea | Formicidae | *Lasius brunneus*<br>(Latreille, 1798) | 50°55'16.0"N 11°34'57.9"E<br>plantain. VR |
| 38 | Hymenoptera | Vespoidea | Vespidae | *Vespa germanica* | 50°55'27.9"N 11°35'00.9"E<br>wild cherry. VR |
| 39 | Hymenoptera | Apoidea | Apidae | *Apis mellifera* | 50°55'24.0"N 11°35'00.6"E<br>Ordinary snowberry. VR |
| 40 | Hymenoptera | Formicoidea | Formicidae | *Lasius* sp. | 50°55'33.3"N 11°33'54.8"E<br>Under dead wood. VR |
| 41 | Hymenoptera | Formicoidea | Formicidae | *Myrmica* sp. | 50°55'34.4"N 11°33'57.0"E<br>Colony under styrofoam. VR |
| 42 | Hymenoptera | Formicoidea | Formicidae | *Camponotus herculeanus*<br>(Linnaeus, 1758) | 50°55'27.2"N 11°35'03.8"E<br>roadside, stone floor. VR |

**Table S6.** Comparison of Hymenoptera at the level superfamily level; data averaged per superfamily (FB = forelimb, ML = middle leg, HL = hind leg).

| Superfamily | Number of individuals N | swing time in s | contact time in s | Duty factor in % | Body length in mm | Speed in mm/s | Stride length in mm | Step width in mm |  |  | Radius of movement, angle in degrees |  |  | Minimum angle (AEP) in degrees |  |  | Maximum angle (PEP) in degrees |  |  |
| --- | --- | --- | --- | --- | --- | --- | --- | --- | --- | --- | --- | --- | --- | --- | --- | --- | --- | --- | --- |
|  |  |  |  |  |  |  |  | FL | ML | HL | FL | ML | HL | FL | ML | HL | FL | ML | HL |
| **Tenthredinoidea** | 2 | 0.06 | 0.25 | 81 | 11.6 | 18.6 | 5.69 | 4.3 | 5.7 | 5.7 | 77 | 56 | 31 | 41 | 69 | 116 | 118 | 124 | 147 |
| **Ichneumonoidea** | 2 | 0.06 | 0.17 | 75 | 7.1 | 20.0 | 4.47 | 2.3 | 3.5 | 3.0 | 75 | 63 | 30 | 32 | 65 | 128 | 107 | 127 | 157 |
| **Cynipoidea** | 1 | 0.05 | 0.13 | 71 | 5.5 | 13.3 | 2.48 | 1.8 | 2.6 | 1.8 | 73 | 52 | 21 | 26 | 71 | 138 | 99 | 123 | 159 |
| **Evanioidea** | 1 | 0.06 | 0.14 | 68 | 12.1 | 23.6 | 4.77 | 2.4 | 3.7 | 3.4 | 82 | 51 | 24 | 25 | 58 | 128 | 107 | 109 | 152 |
| **Vespoidea** | 3 | 0.05 | 0.11 | 69 | 19.4 | 63.2 | 10.73 | 5.2 | 9.9 | 8.2 | 62 | 55 | 31 | 26 | 62 | 118 | 89 | 117 | 149 |
| **Chrysidoidea** | 1 | 0.04 | 0.11 | 75 | 7.6 | 22.0 | 3.49 | 1.9 | 3.7 | 2.5 | 72 | 59 | 29 | 20 | 60 | 132 | 93 | 118 | 161 |
| **Tiphioidea** | 1 | 0.05 | 0.09 | 66 | 12.3 | 42.4 | 6.06 | 2.6 | 4.4 | 4.1 | 67 | 59 | 34 | 29 | 64 | 113 | 96 | 122 | 147 |
| **Apoidea** | 7 | 0.05 | 0.11 | 66 | 13.7 | 43.1 | 7.02 | 4.0 | 6.9 | 5.7 | 69 | 58 | 37 | 30 | 56 | 114 | 99 | 114 | 151 |
| **Formicoidea** | 10 | 0.04 | 0.07 | 63 | 5.95 | 49.4 | 4.18 | 1.6 | 3.0 | 1.9 | 69 | 56 | 25 | 26 | 71 | 138 | 95 | 127 | 163 |
| **overall average** |  | 0.05<br>±0.01 | 0.13<br>±0.05 | 70<br>±5.34 | 10.58<br>±4.24 | 32.8<br>±16.1 | 5.43<br>±2.27 | 2.9<br>±1.2 | 4.8<br>±2.2 | 4.0<br>±2.0 | 72<br>±5.5 | 57<br>±3.5 | 29<br>±4.7 | 28<br>±5.5 | 64<br>±5.2 | 125<br>±9.4 | 100<br>±8.4 | 120<br>±5.7 | 154<br>±5.8 |

**Table S7.** Comparison of Hymenoptera at genus level (v = velocity, BL = body length, FL = front leg, ML = middle leg, HL = hind leg).

| Genus | ID | swing time in<br>s | contact time<br>in s | Duty factor in<br>% | TCS | $v$ to mm/s | $v$ to BL/s | Stride length in<br>mm | Stride length<br>through BL | moving<br>radius,<br>angle in degrees | | | minimal<br>Angle (AEP)<br>in degrees | | | maximum<br>Angle (PEP)<br>in degrees | | |
| --- | --- | --- | --- | --- | --- | --- | --- | --- | --- | --- | --- | --- | --- | --- | --- | --- | --- | --- |
|  |  |  |  |  |  |  |  |  |  | FL | ML | HL | FL | ML | HL | FL | ML | HL |
| *Arge* | 12 | 0.05 | 0.24 | 82 | 0 | 14.3 | 1.3 | 4.38 | 0.39 | 74 | 55 | 25 | 43 | 71 | 116 | 117 | 126 | 142 |
| *Tenthredo* | 11 | 0.06 | 0.26 | 80 | 0 | 22.8 | 1.9 | 7.01 | 0.57 | 80 | 56 | 36 | 39 | 66 | 115 | 118 | 122 | 152 |
| (Cynipidae) | 17 | 0.05 | 0.13 | 71 | 0.13 | 13.3 | 2.4 | 2.48 | 0.45 | 73 | 52 | 21 | 26 | 71 | 138 | 99 | 123 | 159 |
| *Bracon* | 22 | 0.06 | 0.16 | 73 | 0.65 | 15.4 | 3.0 | 3.39 | 0.65 | 69 | 62 | 23 | 29 | 66 | 139 | 99 | 128 | 162 |
| (Ichneumonidae) | 18 | 0.05 | 0.17 | 77 | 0.03 | 24.6 | 2.7 | 5.55 | 0.62 | 82 | 63 | 36 | 34 | 64 | 116 | 115 | 127 | 152 |
| *Gasteruption* | 15 | 0.06 | 0.14 | 68 | 0.10 | 23.6 | 1.9 | 4.77 | 0.39 | 82 | 51 | 24 | 25 | 58 | 128 | 107 | 109 | 152 |
| *Polistes* | 23 | 0.05 | 0.13 | 72 | 0.14 | 55.3 | 3.3 | 10.35 | 0.62 | 62 | 59 | 39 | 31 | 50 | 100 | 93 | 109 | 140 |
| *Vespa* | 06 | 0.04 | 0.10 | 69 | 0.01 | 78.5 | 3.0 | 13.73 | 0.52 | 59 | 54 | 27 | 16 | 71 | 133 | 76 | 126 | 159 |
| *Vespula* | 04 | 0.05 | 0.10 | 65 | 0.26 | 55.8 | 3.6 | 8.10 | 0.53 | 65 | 52 | 26 | 30 | 65 | 122 | 95 | 117 | 148 |
| *Chrysis* | 07 | 0.04 | 0.11 | 75 | 0.13 | 22.0 | 2.9 | 3.49 | 0.46 | 72 | 59 | 29 | 20 | 60 | 132 | 93 | 118 | 161 |
| *Tiphia* | 13 | 0.05 | 0.09 | 66 | 0.66 | 42.4 | 3.5 | 6.06 | 0.49 | 67 | 59 | 34 | 29 | 64 | 113 | 96 | 122 | 147 |
| (Halictidae) | 08 | 0.04 | 0.07 | 62 | 0.47 | 27.0 | 3.9 | 3.02 | 0.44 | 75 | 56 | 37 | 32 | 61 | 113 | 107 | 118 | 150 |
| *Cerceris* | 14 | 0.06 | 0.08 | 57 | 0.65 | 62.2 | 4.6 | 8.90 | 0.66 | 56 | 64 | 37 | 33 | 48 | 115 | 89 | 112 | 152 |
| *Crabro* | 19 | 0.05 | 0.09 | 65 | 0.42 | 41.6 | 3.3 | 5.54 | 0.44 | 71 | 65 | 37 | 20 | 55 | 120 | 91 | 120 | 157 |
| *Andrena* | 5 | 0.05 | 0.12 | 71 | 0.50 | 37.8 | 2.3 | 6.16 | 0.38 | 48 | 51 | 22 | 25 | 55 | 124 | 73 | 106 | 146 |
| *Bombus* | 9 | 0.06 | 0.16 | 72 | 0.19 | 40.5 | 2.1 | 8.78 | 0.45 | 70 | 68 | 39 | 34 | 55 | 123 | 104 | 122 | 162 |
| *Apis* | 3 | 0.05 | 0.12 | 69 | 0.31 | 61.3 | 3.4 | 10.54 | 0.58 | 83 | 55 | 50 | 30 | 52 | 91 | 113 | 107 | 141 |
| *Camponotus* | 42 | 0.05 | 0.17 | 79 | 0.24 | 30.8 | 3.0 | 6.70 | 0.65 | 74 | 65 | 22 | 25 | 69 | 143 | 99 | 135 | 166 |
| *Myrmica* | 29 | 0.05 | 0.09 | 65 | 0.63 | 20.5 | 4.4 | 2.80 | 0.60 | 75 | 61 | 28 | 24 | 81 | 143 | 99 | 142 | 171 |
| *Formica* | 26 | 0.03 | 0.05 | 60 | 0.57 | 61.4 | 10.3 | 4.64 | 0.77 | 68 | 67 | 21 | 23 | 63 | 143 | 91 | 129 | 164 |
| *Lasius* | 37 | 0.02 | 0.03 | 54 | 0.76 | 70.2 | 19.6 | 3.47 | 0.96 | 67 | 55 | 29 | 30 | 70 | 133 | 96 | 125 | 162 |

**Table S8.** Data overview of the Formicidae (FL = front leg, ML = middle leg, HL = hind leg).

| Genus species | ID | Mass (mg) | Swing time (s) | Contact time (s) | Duty factor (%) | Body length (mm) | Speed (mm/s) | Stride length (mm) | Step width (mm) |  |  |
| --- | --- | --- | --- | --- | --- | --- | --- | --- | --- | --- | --- |
|  |  |  |  |  |  |  |  |  | FL | ML | HL |
| *Camponotus* sp. | 42 | 12.3 | 0.05 | 0.17 | 79 | 10.3 | 30.8 | 6.70 | 3.1 | 4.5 | 2.6 |
| *Formica fusca* | 26 | 3.4 | 0.03 | 0.05 | 60 | 6.0 | 61.4 | 4.64 | 1.8 | 3.5 | 1.7 |
| *Myrmica* sp. | 29 | - | 0.05 | 0.09 | 65 | 4.7 | 20.5 | 2.80 | 1.4 | 1.8 | 1.1 |
| *Myrmica* sp | 30 | 3.5 | 0.04 | 0.07 | 61 | 4.8 | 40.7 | 4:11 | 1.6 | 3.4 | 2.3 |
| *Lasius fuliginosus* | 21 | 2.4 | 0.04 | 0.08 | 67 | 5.0 | 26.8 | 3.08 | 1.8 | 3.2 | 2.5 |
| *Lasius niger* (queen) | 27 | 26.1 | 0.04 | 0.05 | 57 | 10.7 | 76.9 | 6.38 | 1.8 | 4.2 | 3.1 |
| *Lasius niger* (worker) | 28 | 1.6 | 0.03 | 0.04 | 59 | 4.4 | 67.2 | 4:17 | 0.9 | 2.5 | 1.4 |
| *Lasius brunneus* | 37 | 1.4 | 0.02 | 0.03 | 54 | 3.6 | 70.2 | 3.47 | 1.1 | 2.2 | 1.4 |
| *Lasius* sp. (infested) | 35 | 2.2 | 0.04 | 0.09 | 69 | 5.7 | 20.0 | 2.62 | 1.6 | 2.8 | 2.1 |
| *Lasius* sp. (uninfested) | 36 |  | 0.03 | 0.04 | 59 | 4.3 | 79.2 | 3.84 | 1.3 | 2.4 | 1.2 |

Among the hymenopteran lineages that we sampled, the genus *Tiphia* was of particular interest as tiphiid females are frequently wingless thus grossly similar to ants, yet are ecologically distinct, spending most of their lives on or under the ground in search of hosts for the brood (Ridsdill Smith 1970; Kimsey 1991). Overall, the sampled *Tiphia* achieved rather average values, except for the comparatively high tripod coordination of 0.66 TCS. Most notably, the *Tiphia* dragged its metasoma while walking, which is a behavior that was never observed in the sampled Formicidae except for trail-laying behavior. This suggests that the locomotion of ants is fundamentally distinct from that of *Tiphia*, despite the ground-based lifestyle of these latter wasps. Given the considerable structural convergence between Tiphiidae and Scoliidae—the sistergroup to Formicoidea + Apoidea—it will be worth investigating the biomechanical architecture and performance of a broader sample of these wasps (see, *e.g.*, Reid 1941; Brothers 1975).

The largest species in our dataset was *Vespa crabro*, with a mass of 190 mg and body length of 2.6 cm. As discussed in the main text, despite having the highest absolute speed of all sampled Hymenoptera (78.5 mm/s), this individual was only slightly faster than the much smaller queen *Lasius niger* that we measured (BL = 10.7 mm; *v* = 76.9 mm/s). Notably, the *Vespa* had a highly irregular gait pattern and, at 0.01, the lowest TCS of all sampled Apocrita. A tripod gait was still observable in the temporal step pattern, although it was extremely delayed within the step cycle (Fig. 16a). Relative to other taxa, the *Vespa* walked in a more meandering fashion rather than straight ahead, thus changing direction and flapping its wings. It is this imperfectly straight walk that may have contributed to the low TCS, as the tetrapod gait may be preferred during direction change or turning (*e.g.*, Zhao *et al.,* 2018). Otherwise, all the prerequisites for a good lap were met for the hornet as the selected measuring chamber offered it enough freedom, and it adhered well to the smooth glass surface of the chamber floor.

Within the Ichneumonoidea we sampled one braconid (*Bracon*) and one ichneumonid (unidentified), which locomoted at a similar speed in relation to their body lengths (3 and 2.7 BL/s). We observed that the ichneumonid shared two unusual behavioral traits with Formicidae to the exclusion of *Bracon*: femorotibial joints raised above the mesosomal dorsum (Fig. 4, Char. 5) and the absence of metasomal dragging during locomotion (Fig. 4, Char. 6). Contrasting with both *Bracon* and the ants, however, the ichneumonid had a very asynchronous gait (0.03 TCS).

In a previous study, Gladun & Gorb (2007) established that the running behavior on a straight surface of an ichneumonid (*Coelichneumon sugillatorius*) was similar to that of an *Arge* (*A. melanochroa*). This is also confirmed by the locomotion results, since the ichneumonid individual showed similar extreme positions of the tarsi (AEP, PEP) and very poor tripod coordination, while the *Arge* individual did not have a tripod gait at all (Table S7).

**Fig. S19.**
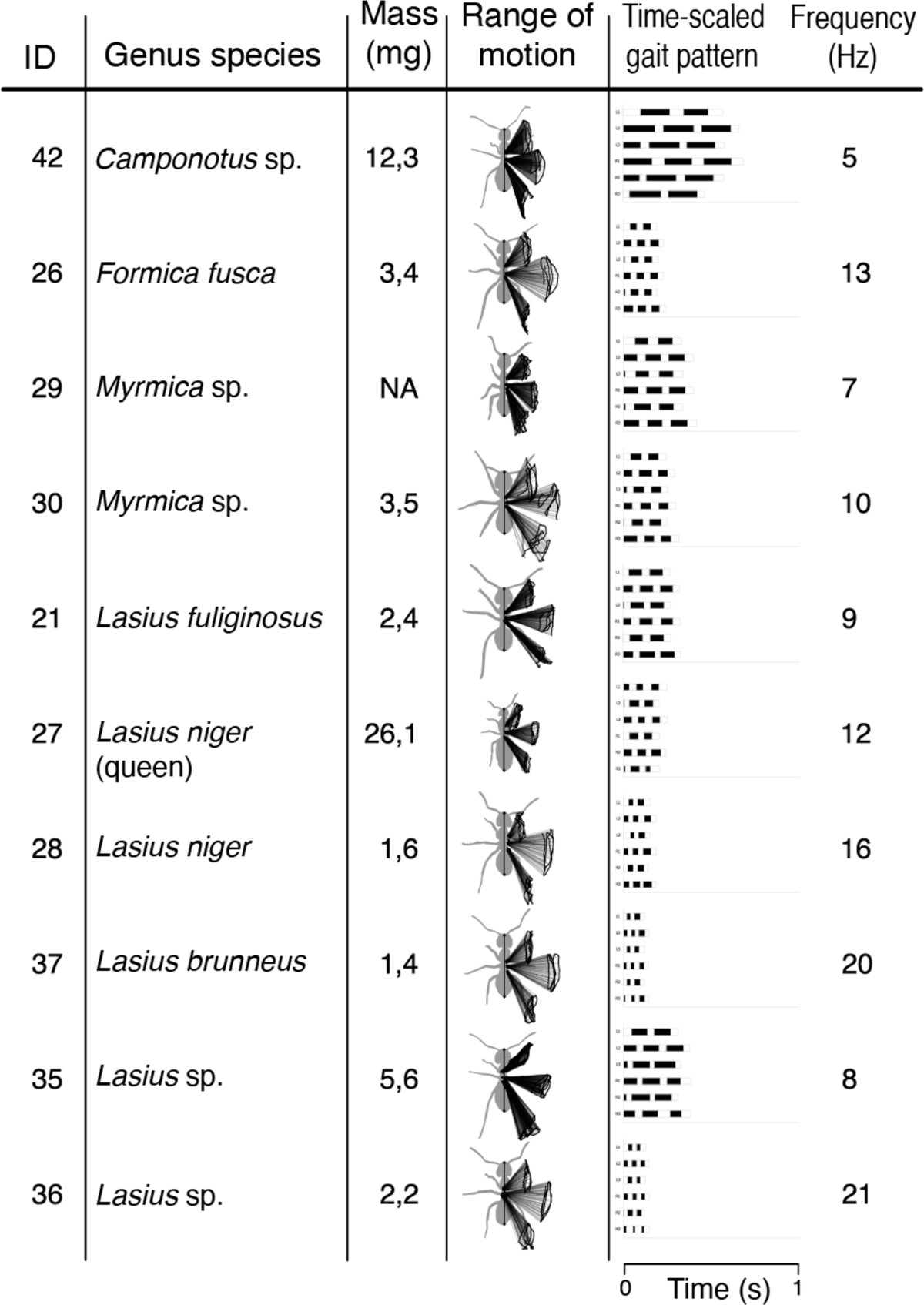
Detail overview of sampled ant locomotory parameters. Rows are arbitrarily ordered.

#### S2.1.3. Results within Formicidae

The sampling of Formicidae in the present study represents the first explicit comparison of temporal and spatial gait patterns of terrestrial locomotion between subfamilies; notably, the swimming mechanics of several subfamilies have been compared by Yanoviak & Frederick (2014) and Schultheiss & Guénard (2021). Regardless, the central results of our ant to non-ant comparisons are robust to subfamily and body size, namely that relative to other Hymenoptera, the sampled Formicidae were able to achieve far greater body-scaled speeds and longer body-scaled strides, and they had larger absolute duty factors and distinct range of motion capacities during straight, flat-surface locomotion (Fig. 1). Although our sampling only included one queen, *Lasius niger* (i27), her performance was very similar to the conspecific worker that we sampled (i28). This consistency indicates that documentation of greater inter-caste, inter-sex, and among-lineage diversity of Formicidae will reveal patterns of locomotory evolution within ants. In the present supplementary section, we will address locomotory patterns with emphasis on *Camponotus*.

The sampled *Camponotus herculeanus* worker was distinct due to its long average swing time and, above all, long contact time; its very high duty factor of 79% clearly exceeded the average 60% of the *Colobopsis schmitzi* (formerly *Camponotus*) from the study by Bohn *et al.,* (2012). At three body lengths per second, our sampled worker only moved at about a third of the speed, and consequently its stride length, which corresponded to about 65% of its body length, was below the relative stride length of *C. schmitzi* at ∼80% (Bohn *et al.,* 2012; Bohn 2007). In addition, the *C. herculeanus* only had a TCS value of 0.24, which was less than half of the other ants examined. On the one hand, this confirmed the observation of Merienne *et al.,* (2020), who found that higher duty factor was associated with lower tripod coordination. On the other, it provides supporting evidence for the thesis that smaller ants walk more stably in the alternating tripod gait than larger ones. The expected reason for this is that large ants have a different gravity centering behavior for the body, which may be caused by the disproportionately large head and has to be compensated for when running (Merienne *et al.,* 2020).

The body length of ∼10.3 mm of the studied *Camponotus* worker was approximately equal to the length of the male *Camponotus fellah* from the work of Tross *et al.,* (2022). That male also achieved a lower tripod coordination and speed than the smaller *C. fellah* ants with 0.5 TCS (Tross *et al.,* 2022). In addition, it could be shown that larger ants leave their legs longer in the contact phase and thus achieve a higher duty factor (Merienne *et al.,* 2020). According to this, both the body size and the body shape of the ant influence its locomotion behavior. Nevertheless, the less efficient running results of the studied *Camponotus* cannot be attributed solely to their size and distinct proportions. The possibilities that the ant was injured or that the measurement chamber could not provide enough space to run at a higher speed must also be considered, as these may result in a longer stride length and higher cadence.

As already mentioned, the other ant genera achieved high TCS values <CAMPO LOW: 0.24; see additional results / discussion> and thus walked in a coordinated and quite synchronous tripod walk. At the top was a *Lasius* individual with a TCS value of 0.76 at a speed of 70.2 mm/s, which is a value comparable to that of the desert ant *Cataglyphis fortis* from Wahl *et al.,* (2015) with 0.77 TCS at a speed of 95.2 mm/s. In contrast, the values of *Formica* (0.57 TCS) and *Myrmica* (0.63 TCS) were similar to the average value of the unloaded *Messor barbarus* of 0.59 TCS (Merienne *et al.,* 2020). Regardless of taxon, these were all relatively high values, especially considering that a completely synchronous alternating tripod gait with a TCS value of 1.0 can almost never be achieved (Weihmann *et al.,* 2015; Wahl *et al.,* 2015).

Similar to *Vespa*, the spatial step pattern of *Myrmica* drifted away from a straight run to change direction. This is probably why it walked at a lower step frequency and a significantly lower speed than the genera *Lasius* and *Formica*, which also exactly reflected the results of Zollikofer (1994b). Since *Myrmica* (i29) escaped from the measurement chamber, it was not possible to use another measurement. The values of the other *Myrmica* (i30) were also out of the question for the genus comparison, as the individual probably suffered from an injured hind leg. However, this is not of great importance for evaluating the results. If one follows the assumption of Zollikofer (1994b), there is no specific locomotion feature that distinguishes the subfamily Myrmicinae, to which *Myrmica* belongs, from the subfamily Formicinae, to which *Lasius*, *Formica*, and *Camponotus* belong.

#### S2.2. Knees and gasters

While extracting the primary variables of interest from our kinematic videos, we observed meaningful behavioral variation with respect to the femorotibial joints and abdomen. Although mentioned in the main text (section 4.2) and elsewhere in the supplementary material (section S2.1.2), these behaviors deserve special attention due to their correlation with the high locomotory speeds of ants (Fig. 4). We specifically observed that the “knee” joints of ants were raised above the mesosomal dorsum, and the abdomen was not dragged except for trail-laying behavior; only a few other sampled taxa displayed these behavioral traits (Table 2). Taken altogether in the phylogenetic-comparative context of the present study, these observations indicate that the static and dynamic posture of ants is distinct relative to other Hymenoptera, which appears to us as likely to be a consequence of their anatomical derivation.

## Notes

### Competing Interest Statement

The authors have declared no competing interest.

